# Unravelling the role of epigenetic regulators during embryonic development of *Rhipicephalus microplus*

**DOI:** 10.1101/2025.07.11.662657

**Authors:** Anderson Mendonça Amarante, Daniel Martins de Oliveira, Marcos Paulo Nicolich Camargo de Souza, Manoel Fonseca-Oliveira, Antonio Galina, Serena Rosignoli, Angélica Fernandes Arcanjo, Bruno Moraes, Alessandro Paiardini, Dante Rotili, Juan Diego de Paula Li Yasumura, Sarah Henaut-Jacobs, Thiago Motta Venancio, Marcelle Uhl, Rodrigo Nunes-da-Fonseca, Luis Fernando Parizi, Itabajara da Silva Vaz Junior, Claudia dos Santos Mermelstein, Lucas Tirloni, Carlos Logullo, Marcelo Rosado Fantappié

## Abstract

Epigenetic modifications are long-lasting changes to the genome that influence a cell’s transcriptional potential, thereby altering its function. These modifications can trigger adaptive responses that impact protein expression and various cellular processes, including differentiation and growth. The primary epigenetic mechanisms identified to date include DNA and RNA methylation, histone modifications, and microRNA-mediated regulation of gene expression. The intricate crosstalk among these mechanisms makes epigenetics a compelling field for the development of novel control strategies, particularly through the use of epigenetic drugs targeting arthropod vectors such as ticks. In this study, we identified the *Rhipicephalus microplus* orthologs of canonical histone-modifying enzymes, along with components of the machinery responsible for m^5^C and ^6^mA-DNA, and m^6^A-RNA methylations. We further characterized their transcriptional profiles and enzymatic activities during embryonic development. To explore the functional consequences of epigenetic regulation in *R. microplus*, we evaluated the effects of various epigenetic inhibitors on the BME26 tick embryonic cell line. Molecular docking simulations were performed to predict the binding mode of these inhibitors to tick enzymes, followed by *in vitro* assessment of their effects on cell viability and morphology. Tick cells exposed to these inhibitors exhibited phenotypic and molecular alterations. Notably, we observed higher levels of DNA methylation in the mitochondrial genome compared to nuclear DNA. Inhibition of DNA methylation using 5’-azacytidine (5’-AZA) was associated with increased activity of the mitochondrial electron transport chain and ATP synthesis, but reduced cellular proliferation. Our findings highlight the importance of epigenetic regulation during tick embryogenesis and suggest that targeting these pathways may offer a novel and promising strategy for tick control.

**Highlights:**

- *R. microplus* presents a complete set of epigenetic enzymes that modify histones, DNA and RNA
- Epigenetic regulation is highly dynamic, and functionally significant during *R. microplus* embryogenesis
- Inhibition of DNA methylation leads to overactivation of the mitochondrial electron transport chain

## INTRODUCTION

The tick *Rhipicephalus microplus* is an obligatory hematophagous ectoparasite that causes major losses to bovine herds. *R. microplus* is a one-host tick and its life cycle consists of the free-living and the parasitic phases^1^. Ticks require a blood meal at each stage (larva, nymph, and adult) to develop into the next stage. Tick eggs do not require external nutrition and are reliant on the yolk provided within the egg for sustenance during development^2^.

The economic losses associated with *R. microplus* parasitism are due to direct effects of the tick itself, which causes skin injuries and long-standing blood loss, leading to anemia and reduction of both weight gain and milk production, or are produced indirectly via transmission of tick-borne pathogens such as *Babesia* spp. and *Anaplasma marginalev*^1,3^. In spite of its huge impact on the economy and human health, current tick control strategies still rely mostly on the use of chemical acaricides, even though selection of resistant tick populations to major used acaricides has been confirmed^4^. This is recognized as a worldwide drawback to successful tick control. Although immunization of cattle against *R. microplus* and other ticks has been recognized as an alternative approach against chemical control strategies, up to this date, an efficient and/or viable vaccine is still lacking^5–7^. Therefore, a deeper understanding of tick physiology is needed as a means to find new molecular targets that can be useful in the development of novel tick control methods.

A promising strategy for controlling gene expression in lower eukaryotes that can cause harm to human and animals has been hypothesized^8–10^. Such a strategy would regulate the expression of genes involved in pathogenicity, differentiation in the life cycle, and other vital aspects of the host and/or pathogen. Therefore, targeting gene regulation mechanisms such as epigenetics would contribute immensely to controlling or even eradicating vectors that cause huge impacts on economy, as well as on human and animal health.

Epigenetic information directs the formation of distinct cellular and organismal phenotypes from a common genome. For example, the ability of insects to develop phenotypes appropriate to their environment relies on epigenetic information^11–15^. Epigenetics is specifically concerned with heritable changes to gene regulation that occur in response to intercellular and extracellular environmental cues. Broadly defined, epigenetic information can take many forms, since factors at many levels can stably affect gene regulation. Thus, the field of molecular epigenetics is generally concerned with molecular mechanisms that directly affect, alter, or interact with chromatin.

Not only in mammals, but also in arthropods, epigenetic modifications play significant roles in the control of multiple biological processes that regulate development, growth, reproduction, behavior, immunity and castle differentiation^16–21^.

DNA methylation is perhaps the most faithfully heritable form of epigenetic information. Unlike other forms of epigenetic information, however, DNA methylation is essentially absent from the genomes of several insect groups, including the dipterans *Drosophila* and *Aedes aegypti*^22,23^. In recent years, DNA methylation has nevertheless captured the attention of entomologists, driven in large part by a desire to understand the importance of DNA methylation to developmental plasticity^24–27^.

The majority of DNA in the metazoan nucleus is incorporated into nucleosomes, which are composed of approximately 147 bp of DNA wrapped around a protein complex composed of eight histone proteins^28,29^. Therefore, many regulatory processes in eukaryotes have been linked to the alteration of histone–DNA interaction. There are several important ways in which nucleosomes can be altered to impact the regulation of genes. Histone posttranslational modifications (hPTMs) are a diverse set of epigenetic signals that typically occur on histone protein N-terminal amino acid tails^28,29^. There are several ways in which hPTMs can alter transcription. First, the association between the target histone and underlying DNA can be directly impacted by the addition of an acetyl or methyl groups to a histone protein, which may consequently increase or decrease the ability of transcription factors to access DNA28,29.

Histone proteins have been studied extensively in *Drosophila*^17,30^, but only recently have histones been investigated in non-model insect taxa^31–33^. Histone protein modifications are functionally conserved in essentially all eukaryotes, including the arthropods^31–33^. Thus, in arthropods, histone proteins may be directly implicated in the mediation of phenotypic plasticity^34^.

N^6^-methyladenosine (m^6^A) is the most abundant internal epigenetic modification in eukaryotic mRNA, which has profound effects on RNA metabolism, including RNA stability^35^, splicing^36^, translation^37^ and RNA-protein interactions^38^. In the last decade, numerous studies revealed that m^6^A modification plays a vital role in regulating eukaryotic growth and development^39^, cell differentiation^40^, reproduction^41^, DNA-damage response ^42^, circadian rhythms^43^ and cancer induction^44^.

The importance of epigenetic modifications during mammalian embryogenesis is well described and characterized^45^. However, the role of epigenetics in arthropod embryogenesis, with the exception of *Drosophila*, is far from being comprehended^46–48^. In this regard, the role of epigenetics on the biology of an important blood-feeding ectoparasite group of arthropods, the ticks, is still scarce^49–51^ and thus, deserves attention.

In the present paper, we aimed at characterizing the role of epigenetic modifications in *Rhipicephalus microplus* during embryogenesis, thus providing a new layer of knowledge for tick biology, as well as providing novel molecular targets for tick control.

## METHODS

### Tick Maintenance

Ticks of the species *R. microplus* (Porto Alegre strain) were obtained from a colony at the Universidade Federal do Rio Grande do Sul, Brazil in accordance with previously described procedures^52^. Animals used in the experiments were housed at Faculdade de Veterinária, Universidade Federal do Rio Grande do Sul (UFRGS) facilities. This research was conducted according to the ethics and methodological guidance, in agreement with the International and National Directives and Norms for Animal Experimentation Ethics Committee of Universidade Federal do Rio Grande do Sul (process number 45209). These ticks, which were free of *Babesia* spp., were maintained on calves in an area that lacked other tick species. Naturally detached fully engorged females. To ensure appropriate egg collection, the females were fixed in metal supports with tape and maintained in an incubator at a constant temperature of 28°C and humidity level of 80%. Oviposition began several days after the females were collected. During oviposition, eggs were collected daily and kept in petri dishes within the same incubator. This collection scheme was utilized to obtain eggs over the course of 21 days, as follows: days 6, 9, 12, 15, 18 and 21. Embryonic fixation and DAPI staining were carried out exactly as previously described^53^.

### Genome-wide identification of epigenetic enzymes in *R. microplus*

We obtained the latest *R. microplus* functional annotations (BIME_Rmic_1.3) from NCBI RefSeq *R. microplus* genome assembly BIME_Rmic_1.3 - NCBI – NLM). The epigenetic enzymes from *R. microplus* were identifyed after performing a BlastP (e-value ≤ 1e-10, identity ≥ 25% and query coverage ≥ 50%)^54,55^, using sequences of *Ixodes scapularis* epigenetic enzymes against *R. microplus* transcriptome. *Drosophila* epigenetic enzyme homologs were also blasted against *R. microplus* database. The epigenetic enzymes identified in *R. microplus* were double-checked by aligning them with the human homologs. The conserved domain architectures were rendered with DOG (Domain Graphs, Version 1.0)^56^. Multiple sequence alignments were performed using Clustalw^57^.

### Phylogenetic analysis

We assembled a non-redundant genome dataset from diverse arthropods, capturing key evolutionary relationships. Complete genome sequences were retrieved from GenBank, and target protein sequences from *R. microplus* were extracted and saved as .faa files. To identify orthologs, we used OrthoFinder (v3.0.1b1), followed by redundancy filtering with Vsearch (v2.30.0), removing sequences with >99% identity. Multiple sequence alignment was performed using MAFFT (v7.525) with 1000 iterations, and phylogenetic trees were built with IQ-TREE2 (v2.3.6) using 1000 bootstrap replicates. Protein domain annotations were obtained using Hmmscan (HMMER v3.4) and the Pfam database, with domains filtered based on an E-value threshold of 10⁻³. Finally, data visualization was performed in R using ggplot2 (v3.5.1).

### Molecular docking

Amino acid sequences of target enzymes were obtained from UniProt^59^. Sequences were submitted to the AlphaFold3 server^60^ with default parameters (5 recycles, template mode disabled, AMBER relaxation). Predicted structures were ranked by per-residue confidence scores (pLDDT > 80). Predicted alignment error (PAE) maps confirmed domain stability (PAE < 10 Å for intra-domain residues). Models were refined via energy minimization in GROMACS (2023.1; steepest descent, 5000 steps; and protonated at pH 7.0, and charges assigned via AMBER ff14SB force field. Grid boxes for docking were defined using residues homologous to human enzyme active sites. Human METTL3/METTL14 heterodimer, DNA methyltransferase, and histone deacetylases were retrieved from the RCSB Protein Data Bank^61^. Polar hydrogens and Kollman charges were added using AutoDockTools (ADT v1.5.7). For docking with STM2457 (Mettl3/Mettl14 inhibitor), 5-azacytidine (5’-AZA; DNMT inhibitor), and trichostatin A (TSA; HDAC inhibitor), co-crystallized water molecules and non-essential ions were ignored. Structures were converted to PDBQT format to define atomic partial charges and rotatable bonds. Molecular docking was performed using AutoDock Vina^62^, integrated into DockingPie^63,64^ within PyMOL. Exhaustiveness was set to 32 for Vina, and Lamarckian genetic algorithm parameters (256 runs, 25 million energy evaluations) were used for AutoDock. The top 10 poses per ligand were clustered by root-mean-square deviation (RMSD < 2.0 Å) and ranked by binding affinity (ΔG, kcal/mol). Visual analysis of hydrogen bonding, hydrophobic interactions, and steric complementarity was performed in PyMOL.

### Nucleic acids isolation and quantitative real-time PCR

Genomic DNA and Total RNA were isolated from 50mg of tick eggs of the different developmental stages (days 6 through 21 after egg hatching). Genomic DNA was purified using the DNeasy Blood & Tissue kit (Qiagen) following the manufacturer’s instructions. Total RNA was obtained using the RiboPure kit (Ambion) followed by DNase treatment (Ambion) and cDNA synthesis (SuperScript III First-Strand Synthesis System, Invitrogen), following the manufacturer’s instructions. Quantitative reverse transcription gene amplifications (qRT-PCR) were performed with StepOnePlus Real-Time PCR System (Applied Biosystems) using the Power SYBR Green PCR Master Mix (Applied Biosystems). The comparative *Ct* method was used to compare mRNA abundance. In all qRT-PCR analyses, the *R. microplus* elongation factor 1α (*ELF1α*) gene was used as an endogenous control gene^58^. All oligonucleotide sequences used in qRT-PCR assays are listed in the Table S1.

### Dot blot

Two hundred nanograms of genomic DNA or total RNA were spotted on a polyvinylidene fluoride membrane (PVDF, Roche). The nucleic acids were fixed by baking the membranes at 80°C for 1 hour. The membranes were then blocked in the blocking buffer (5% skim milk in PBST) for 2 h at room temperature, and incubated with anti m^5^C or m^6^A monoclonal antibodies (Table S2) overnight at 4 °C. After three washes with wash buffer (5% skim milk in PBST) for 10 min each, the membrane was incubated with an anti-secondary antibody (Table 1) for 60 min at room temperature. After three washes as above, signals were detected by chemiluminescence using the SuperSignal West Dura (Thermofisher), and the Amersham Image 600 (GE HealthCare) Imaging System. Nucleic acid input controls were stained with 0.04% methylene blue.

**Table 1.** List of antibodies used in this study.

| **Primary antibodies** | **Code** |
| --- | --- |
| Anti-H3, Cell Signaling | 14269 |
| Anti-H3K27ac, Cell Signaling | 8173 |
| Anti-H3K14ac, Cell Signaling | 7627 |
| Anti-H3K4me3, Cell Signaling | 9727 |
| Anti-H3K9me2, Cell Signaling | 4658 |
| Anti-H3K9me3, Cell Signaling | 13969 |
| Anti-H3K27me3, Cell Signaling | 9733 |
| Anti-m6A, Invitrogen | MA5-35350 |
| Anti-m5C, Abcam | ab214727 |
| Anti-NDUFS3, Abcam | ab14711 |
| **Secondary antibodies** | **Code** |
| Anti-mouse, Invitrogen | 31430 |
| Anti-Rabbit, Invitrogen | 32460 |
| Anti-Rabbit Alexa fluor 488, Invitrogen | A11001 |
| Anti-Rabbit Alexa fluor 568, Invitrogen | A11004 |

### Western blot

Protein extracts were prepared as previously described^20^. Briefly, total protein extracts from 50mg of tick eggs of the different developmental stages (days 6 through 21 after egg hatching), were carried out by homogenization in TBS containing a protease inhibitor cocktail (Sigma). Proteins were recovered from the supernatant by centrifugation at 14.000xg, for 15 min. at 4°C. Protein concentration was determined by the Bradford Protein Assay (Bio-Rad). Western blots were carried out using the secondary antibodies (Table S2) with a 1:5.000 dilution. The primary epigenetic monoclonal antibodies (ChIP grade) used were anti-H3K9ac, anti-H3K27ac, anti-H3K27me^3^, anti-H3K4me^3^, anti-H3K9me^2^, and anti-H3K9me^3^ (Table 2), according to the manufacture’s instructions. For all antibodies, a 1:4.000 dilution was used. For normalization of the signals across the samples, an anti-histone H3 antibody (Table S2) was used.

**Table 2.** Protein sequence homology of epigenetic regulators across *Rhipicephalus microplus*, *Bos taurus*, and *Homo sapiens*. GenBank accession numbers are listed for each protein, along with the percentage identity of full-length sequences between *R. microplus* and the other species (*B. taurus* and *H. sapiens*). Sequence alignments were performed using [*tool/algorithm*, e.g., BLASTP] with default parameters.

| Protein name | *R. microplus* | *B. taurus* |  | *H. sapiens* |  |
| --- | --- | --- | --- | --- | --- |
|  | GenBank ID | GenBank ID | Identity | GenBank ID | Identity |
| CBP/P300 | XP_037286635.1 | NP_001157494.1 | 61% | XP_016878433.1 | 61% |
| EZH2 | XP_037288170.1 | XP_059741477. | 59% | XP_005250020.1 | 58% |
| EHMT | XP_037291703.1 | XP_024854576.1 | 52% | XP_054219807.1 | 50% |
| SETDB1-A | XP_037277427.1 | NP_001178317.1 | 46% | NP_001380890.1 | 45% |
| SETD2 | XP_037282969.1 | XP_024849516.1 | 49% | NP_001035889.1 | 49% |
| SETD4 | XP_037281861.1 | NP_001192753.1 | 32% | NP_001273681.1 | 34% |
| SETD7 | XP_037279979.1 | XP_024833097.1 | 47% | NP_001311433.1 | 52% |
| GCN5 | XP_037291980.1 | XP_024853358.2 | 63% | XP_006721881.1 | 60% |
| HDAC1 | XP_037271577.1 | NP_001032521.1 | 87% | NP_004955.2 | 85% |
| HDAC4 | XP_037291540.1 | XP_059741238.1 | 47% | AAH39904.1 | 47% |
| HDAC6 | XP_037286423.1 | XP_024843702.1 | 45% | AAH69243.1 | 47% |
| DNMT1 | XP_037279276.1 | XP_024849600.1 | 57% | NP_001124295.1 | 57% |
| DNMT3A | XP_037270226.1 | XP_059747024.1 | 39% | NP_715640.2 | 39% |
| DNMT3B | XP_037270264.1 | XP_024856107.1 | 47% | NP_001411281.1 | 47% |
| N6AMT1 | XP_037277348.1 | NP_001076982.1 | 48% | NP_037372.4 | 50% |
| METLL3 | XP_037282032.1 | XP_024853529.1 | 50% | NP_062826.2 | 54% |
| METLL14 | XP_037281356.1 | NP_001077183.1 | 64% | NP_066012.1 | 64% |

**Table 3.**
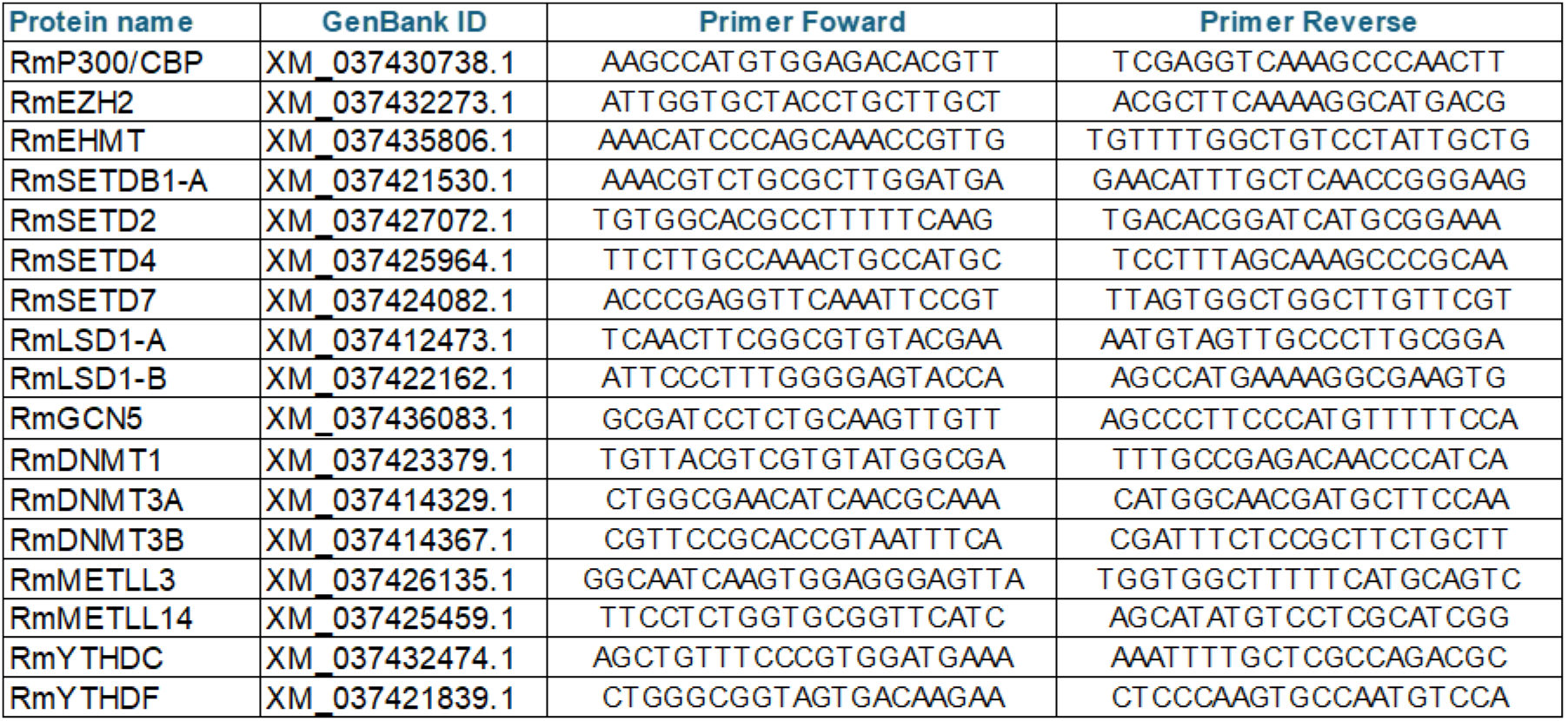
List of primer sequences used in this study. GenBank identification is provided for all genes.

### Statistical analysis

Densitometry analyses were performed using ImageJ software IJ1.46r. For the colocalization analysis, the Coloc2 plugin from ImageJ IJ1.46r was used. All statistical analyses were performed with the GraphPad Prism statistical software package (Prism version 8.4, GraphPad Software, Inc., La Jolla, CA). Asterisks indicate significant differences (*, p < 0.05; **, p < 0.01; ***, p < 0.001; ns, not significant).

### Cell culture and cell viability assay

The BME26 tick embryo cell line was originally obtained as previously described^65^ and maintained according to the established protocols^65^. Cells were maintained in Leibovitz L-15 medium (Sigma-Aldrich®), supplemented with amino acids, glucose, mineral salts, and vitamins, as previously described^65^. The medium was diluted in sterile water (3:1), followed by addition of 10% tryptose phosphate broth (Sigma-Aldrich, #T8782), 10% fetal calf serum (Nutricell®, inactivated by heating), and commercial antibiotic Penicillin-Streptomycin (Gibco, #15140122) was diluted in the medium (1:100), according to the manufacturer’s instructions. Cells (4.1 x 10^5^) were incubated on 24-well multiplates with 25, 50 or 100μM of the epigenetic inhibitors STM2457^66^, 5-Aza-2’-deoxycytidine (Sigma-Aldrich), and Trichostatin A (Sigma-Aldrich) for 48h. To count cells using a Neubauer chamber, a cell suspension is placed on the grid and cells are counted under a light microscope. Cell viability assay was performed using the CellTiter-Glo® Luminescent Cell Viability Assay” (Promega).

### Fluorescence Microscopy

A total of 4.1 x 10^5^ BME26 cells were cultured on glass coverslips placed in a 24-well plate. Cells were fixed with 4% paraformaldehyde for 1 hour and subsequently permeabilized for 15 minutes in 1X PBS containing 0.15% Triton X-100. Following permeabilization, cells were incubated in blocking buffer (1X PBS, 0.05% Tween-20, and 10% albumin) for 1 hour at room temperature. Cells were then incubated overnight (16h) at 4°C with the appropriate primary antibodies (as listed in Table S2), diluted 1:5000 in blocking buffer. After primary antibody incubation, cells were washed three times for 10 minutes each in 1X PBS containing 0.05% Tween-20. Subsequently, cells were incubated for 1 hour with an Alexa Fluor 488-conjugated anti-rabbit secondary antibody (Table S2), diluted 1:1000 in blocking buffer. Afterward, cells were washed again three times for 10 minutes each. Coverslips were mounted onto glass slides using 10μL of ProLong™ Gold Antifade Mountant with DAPI (Thermo Fisher). Immunofluorescence images were acquired using a Zeiss LSM 710 confocal microscope.

For Phalloidin Staining, BME26 cells were cultured on glass coverslips in a 24-well plate. They were then fixed with 4% paraformaldehyde for 1 hour and permeabilized for 15 minutes in 1X PBS with 0.15% Triton. To reduce background from nonspecific staining, the cells were incubated with 1% BSA in PBS for 40 minutes at room temperature. For actin staining, the cells were incubated with Rhodamine Phalloidin reagent for 40 minutes at room temperature. Subsequently, the cells were washed with PBS three times for 10 minutes each. The coverslips were then mounted onto glass slides with 10 µL of ProLong Gold Antifade Mountant with DNA Stain DAPI (Thermo Fisher). Immunofluorescence images were acquired using a Zeiss LSM710 confocal microscope.

### Respirometry

High-resolution respirometry was performed using an Oroboros Oxygraph-2k at 32°C with a suspension of 2 × 10⁶ cells/mL in culture medium without fetal bovine serum (FBS). Basal oxygen consumption was recorded until signal stabilization. Subsequently, oligomycin (0.2 µg/mL) was added to inhibit ATP synthase and assess proton leak respiration, followed by stepwise titration of carbonyl cyanide-4-(trifluoromethoxy)phenylhydrazone (FCCP) to determine maximal uncoupled respiration and spare respiratory capacity. Finally, antimycin A (2 µM) was added to inhibit complex III and quantify non-mitochondrial respiration. Oxygen consumption rates were analyzed using DatLab software and corrected for residual (non-mitochondrial) respiration. ATP-linked respiration was calculated as the difference between basal respiration and proton leak, and spare respiratory capacity as the difference between maximal uncoupled and basal respiration. All experiments were performed in three biological replicates. Data are presented as mean ± SEM, and statistical analysis was performed using two-way ANOVA.

## RESULTS

### The epigenetic machinery is encoded within the *R. microplus* genome

Given the limited number of epigenetic studies in ticks and the vast diversity of histone modifications in eukaryotic chromatin, we chose to investigate the presence of epigenetic enzyme homologs in the *R. microplus* genome (Figure 1 and Table S3). For histone-modifying enzymes, we focused on two major histone acetyltransferases (HATs), CBP/p300 and GCN5, along with six histone methyltransferases (HMTs): EZH2, EHMT, SETDB, SETD2, SETD4, and SETD7. Additionally, we examined three histone deacetylases, HDAC1, HDAC4, and HDAC8, representing Class I, II, and III, respectively^67^. We also identified the complete set of enzymes responsible for 5-methylcytosine (m^5^C) DNA methylation, including DNMT1, DNMT3A, and DNMT3B, as well as components involved in N^6^-methyladenosine (m^6^A) RNA methylation: the writer proteins METTL3, METTL14, and WTAP, and the reader proteins YTHDC and YTHDF. Importantly, the roles of these epigenetic enzymes in embryonic development of other organisms have been well documented^69–73^.

**Figure 1.**
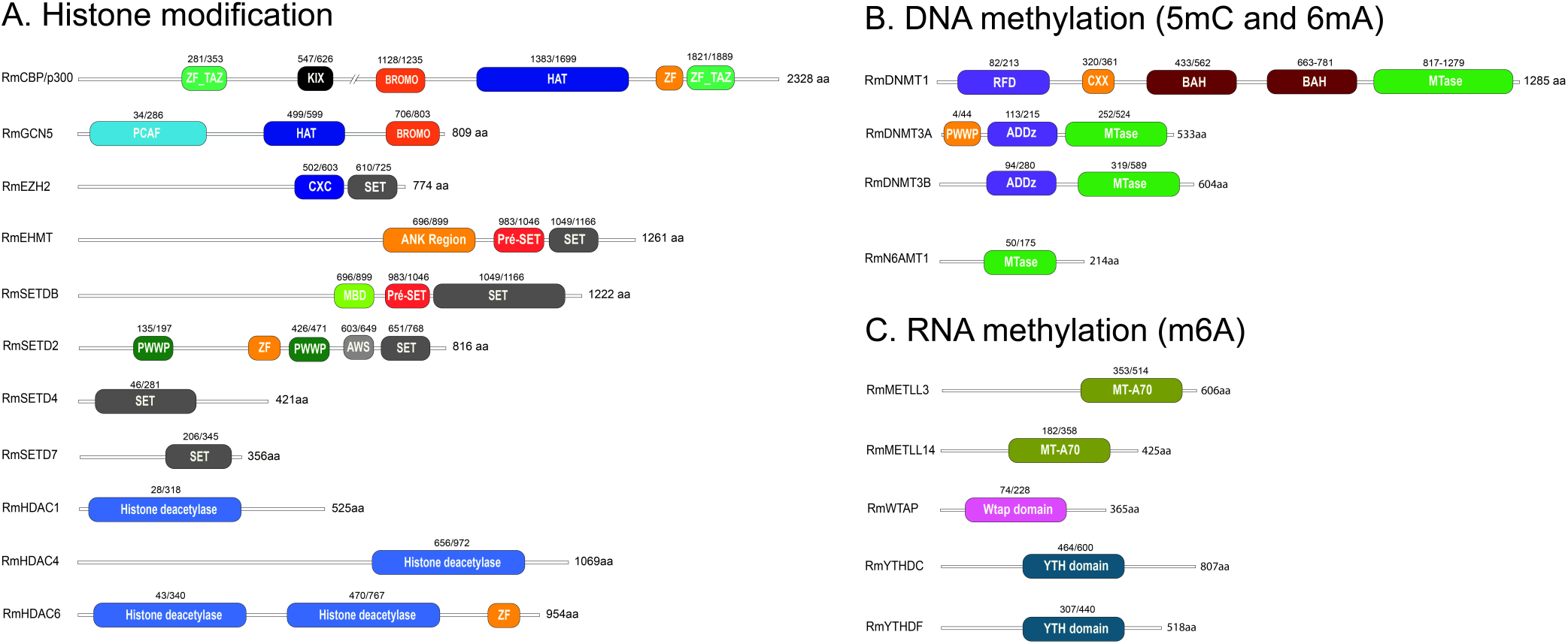
Conserved domains in epigenetic regulators of *Rhipicephalus microplus*. The domain architecture of epigenetic regulators in *R. microplus* is shown in A-C, with gene accession numbers listed in Supplemental Table 3. A comparative analysis of full-length amino acid sequences, including functional domain annotations, between *R. microplus* epigenetic regulators and their mammalian counterparts is provided in Figure S1, A-Q).

Analysis of the sequence alignments for all proteins (Figures S1A–Q), along with the percentage identity data (Table S1), reveals a high level of conservation within the catalytic domains between tick and mammalian homologues.

The phylogenetic and domain architecture analysis of *R. microplus* Ezh2 orthologs in related arthropods revealed distinct evolutionary patterns and conserved functional domains (Figure S2A). OrthoFinder identified 45 orthologs across 20 species, including *Apis*, *Solenopsis*, and *Drosophila*. Phylogenetic reconstruction with IQ-TREE2 supported strong clustering of *R. microplus* with *Ixodes scapularis* (XP_029824349.2, XP_029824350.2), reflecting their shared taxonomic order (Parasitiformes), with bootstrap values >90%. Domain annotation via HMMER/Pfam confirmed conserved functional motifs: SET, preSET_CXC, PRC2_HTH_1, and EZH2_MCSS domains were universally present, while FLYWCH and Vfa1 domains were lineage-specific.

Phylogenetic and domain architecture analysis of *R. microplus* CBP/p300 orthologs (Figure S2B) revealed conserved functional domains and evolutionary relationships. Phylogenetic reconstruction via IQ-TREE2 demonstrated strong clustering of *R. microplus* with arachnid species such as *Parasteatoda tepidariorum* (XP_015910461.1, XP_019822458.1) (bootstrap >85%), reflecting shared chelicerate ancestry. Domain annotation using HMMER/Pfam confirmed the universal presence of most domains, with only KIX and KIX_2 being alternated in certain taxa. Multiple sequence alignment (MAFFT) highlighted divergent regions in the Creb-binding domain of *R. microplus* and its cluster, suggesting functional divergence unique to acarines.

### Embryonic development of *R. microplus*

To investigate epigenetic dynamics during *R. microplus* embryogenesis, we reproduced a previously established protocol that defines embryological stages using DAPI staining^53^, covering the developmental window from day 6 to day 21 (Figure 2, showing both lateral and dorsal views). Although the initial cleavages occur between days 1 and 3 marked by early cell divisions and migration toward the yolk periphery, we chose not to include these stages in our study. This decision was based on the low number of cells and nuclei present at these time points, as indicated by the minimal detection of histone molecules (Figure S3). The end of days 5 and 6 corresponds to stage 7 of development^53^, during which the germ band is visible along with defined head, thoracic segments, and a posterior growth zone. Segmental grooves begin to emerge at this stage. By day 9 (stage 10), the first three pairs of legs (L1–L3) elongate and reach the ventral midline, while the fourth leg (L4) remains undeveloped. Leg segmentation becomes evident, and the germ band begins to retract. At day 12 (stage 11), the posterior region continues retracting and migrates ventrally in an inverted orientation. The legs shift to a lateral position, and the embryo begins to appear more condensed. Leg segmentation becomes more pronounced in L1–L3, while L4 remains minimal. The chelicerae and pedipalps also relocate to the ventral side. Between days 15 and 18 (stage 13), dorsal closure nears completion, and the prosoma reaches its final anatomical position. From days 18 to 21 (stage 14), the larva is fully formed and prepares to hatch from the egg.

**Figure 2.**
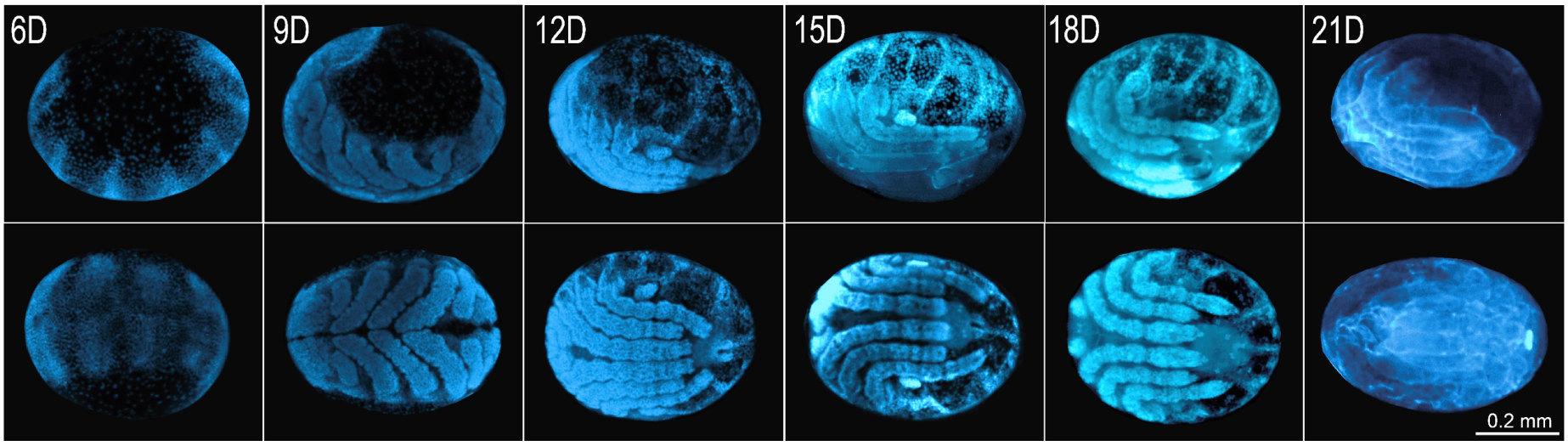
Embryonic development of *Rhipicephalus microplus*. Developmental stages were visualized using DAPI staining to highlight nuclear morphology. The stages, previously described in the literature^53^, are categorized from stage 1 to stage 14, corresponding to days 0 through 21 of embryogenesis. Due to the lack of enough cells in eggs in the initial stages of development, we conducted experiments using eggs from day 6 (stage 7) to day 21 (stage14). Upper panel: lateral view of the eggs. Bottom panel: dorsal view of the eggs.

### Transcriptional profile of epigenetic proteins throughout *R. microplus* embryonic development

We assessed the transcript levels of genes encoding enzymes involved in histone, DNA, and RNA modifications in *R. microplus* embryos (Figure 3A–C). In Figure 3A, the two histone acetyltransferases (HATs), *CBP/p300* showed elevated expression from day 6 to day 15, followed by a marked decline between days 18 and 21. In contrast, *Gcn5* expression increased during the later stages of development, from day 15 to 21. Among the histone methyltransferases (HMTs), *EZH2*, *EHMT*, and *SETD4* maintained relatively constant expression throughout embryogenesis (days 6– 21). In contrast, *SETDB* and *SETD2* exhibited higher transcript levels during early development (days 6–15), while *SETD7* displayed a distinct pattern: elevated expression from day 6 to 12, a near-complete absence of transcripts from day 15 to 18, followed by a significant reappearance of transcripts at day 21.

**Figure 3.**
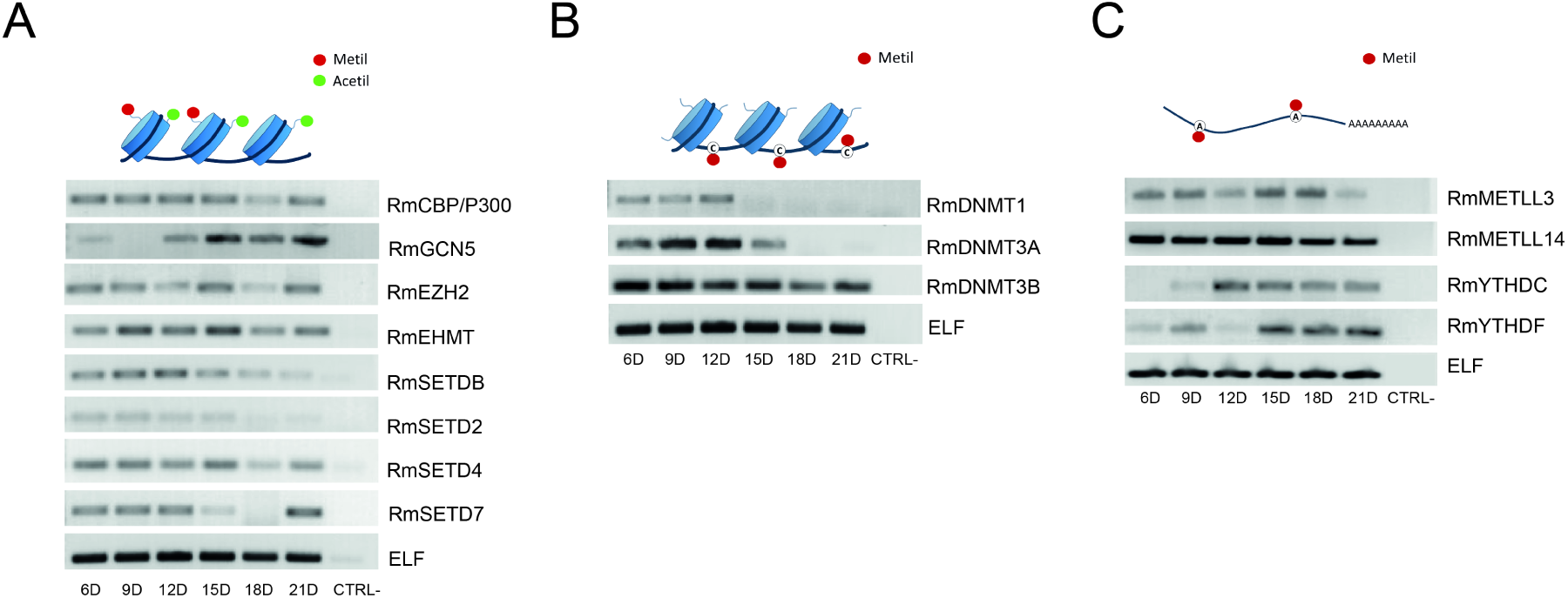
Transcriptional profile of *Rhipicephalus microplus* epigenetic regulators during embryonic development. Enzymes involved in histone modifications (A) were selected based on their well-documented roles in eukaryotic embryogenesis^45^. For m^5^C DNA methylation (B), expression levels of both *de novo* and maintenance DNA methyltransferases were analyzed. For m^6^A RNA methylation (C), the expression of key writer and reader enzymes was evaluated. Elongation factor 1α (ELF) was used as the housekeeping gene for normalization^58^. RT-PCR was performed to assess gene expression, and the amplified products were visualized by agarose gel electrophoresis. Each RT-PCR reaction was independently repeated a minimum of six times to ensure reproducibility.

For the DNA methylation enzymes (Figure 3B), *DNMT1* and *DNMT3a* expression appeared to be restricted to the early stages of development (days 6 to 12), with *DNMT1* showing notably lower transcript levels compared to *DNMT3a*. In contrast, *DNMT3b* exhibited high expression levels throughout the entire developmental period.

Regarding the m^6^A RNA methylation machinery (Figure 3C), the genes encoding the two writer enzymes, *METTL3* and *METTL14*, exhibited relatively constant expression throughout embryogenesis (days 6–21), with *METTL14* showing significantly higher transcript levels than *METTL3*. Among the genes encoding m^6^A reader proteins, *YTHDC* transcription began at day 12 and remained steady through day 21. In contrast, *YTHDF* showed very low expression at days 6 and 9, was undetectable at day 12, and reappeared from day 15 to 21.

### Activity of the epigenetic enzymes throughout *R. microplus* embryonic development

Notably, we observed significant modulation of histone modifications during tick embryogenesis (Figure 4A–F). We examined three histone marks associated with gene activation: H3K27ac, catalyzed by CBP/p300; H3K14ac, catalyzed by Gcn5; and H3K4me^3^, catalyzed by SETD2, SETD4, and/or SETD7 (Figure 4A–C). CBP/p300 activity was nearly undetectable on day 6 of embryonic development but peaked on days 9 and 12, followed by a gradual decline from day 15 to day 21 (Figure 4A). In contrast, Gcn5 displayed an inverse pattern, with minimal acetylation on days 6 and 9, and a progressive increase in activity from day 12 through day 21 ( Figure 4B). Similarly, the activity of SETD2/4/7 was weak on day 6 but increased markedly from day 9 to day 21 (Figure 4C). Regarding repressive histone marks, the activities of EZH2, SETDB, and EHMT were undetectable on days 6 and 9, but increased significantly from day 12 to day 21 (Figure 4D–F). Notably, EzH2 and EHMT showed particularly high activity during the later stages of embryogenesis (Figure 4D and 4F).

**Figure 4.**
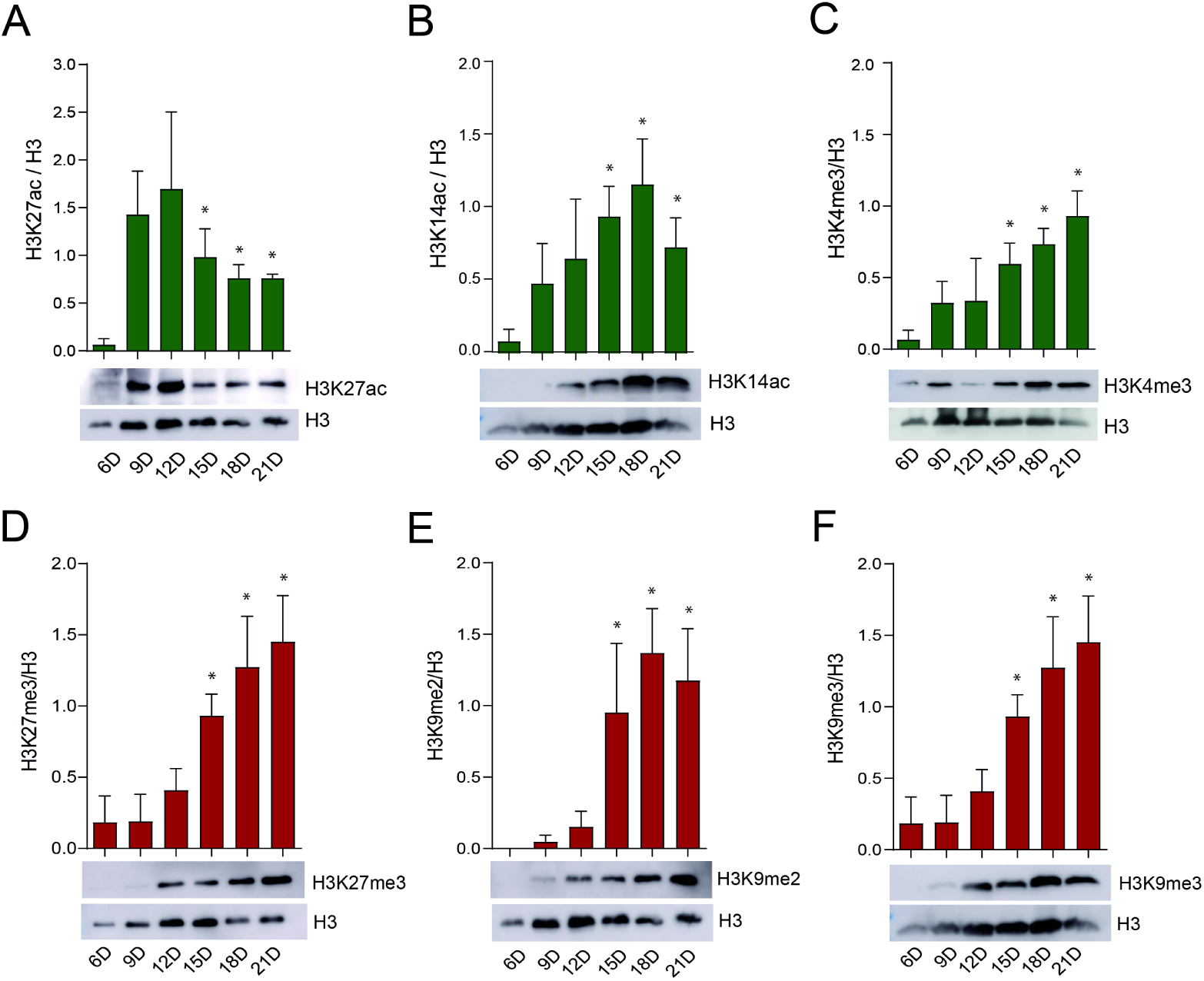
Catalytic activity of *Rhipicephalus microplus* histone-modifying enzymes during embryonic development. Chromatin activation (A-C, green bars) and repression (D-E, red bars) were assessed by Western blot analysis using monoclonal antibodies. Western blotting was performed on six independent biological replicates; a representative blot is shown. Histone H3 was used as a loading control. The intensity of the bands was quantified by densitometry analysis plotted as a graph using ImageJ (NIH Software). Error bars represent the standard error of the mean (SEM). Statistical significance was determined using Student’s *t*-test. *p* < 0.05 is considered significant (*).

Regarding nucleic acid modifications, significant changes were observed only in mRNA m^6^A methylation (Figure 5A). Specifically, METTL3/METTL14 activity was high on days 6 and 9, followed by a marked and gradual decline from day 12 to day 21 (Figure 5A). In contrast, ^6^mA and m^5^C DNA methylation levels remained relatively stable throughout embryonic development (Figure 5B and C, respectively).

**Figure 5.**
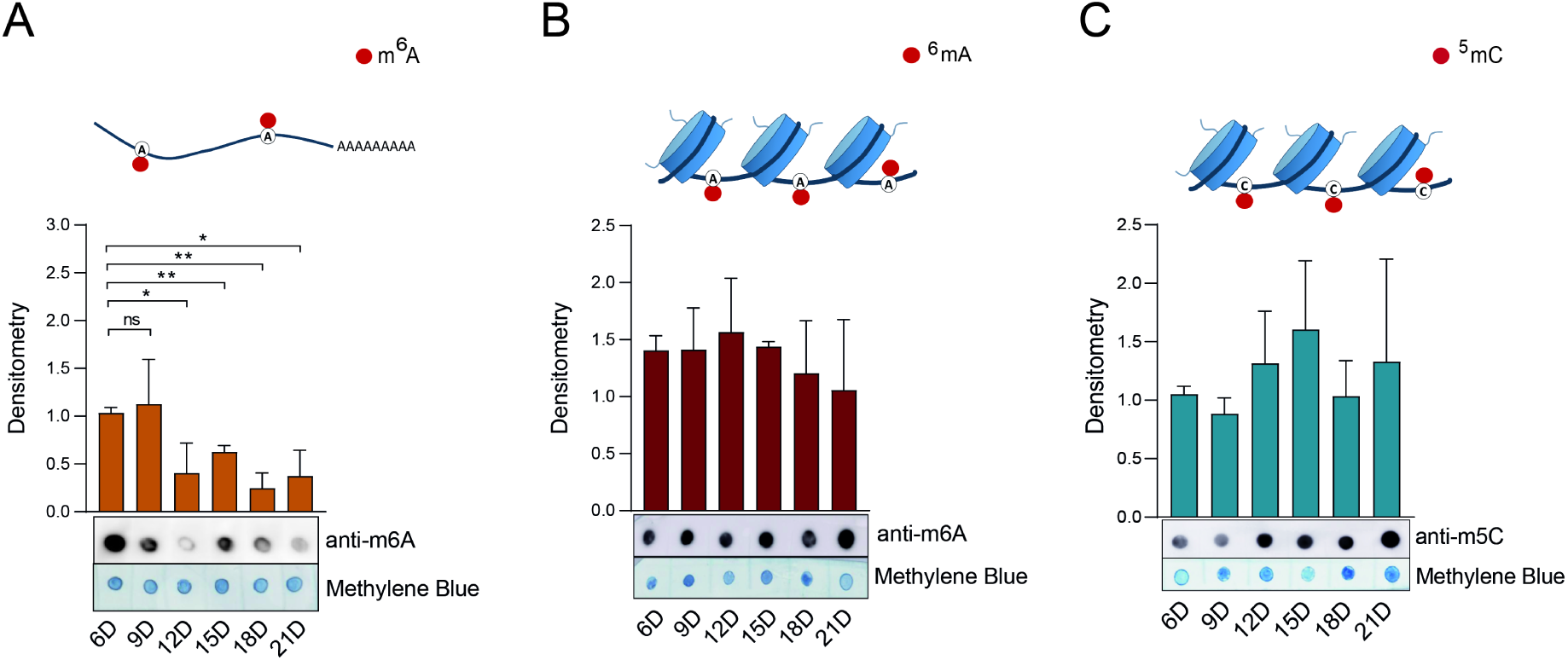
Catalytic activity of *Rhipicephalus microplus* m^6^A-DNA/RNA (A,B) and m^5^C-DNA (C) methyltransferases during embryonic development. Dot blot analyses were performed using six independent biological replicates; one representative blot is shown. Nucleic acids were stained with methylene blue and used as a loading control. Band intensities were quantified by densitometric analysis using ImageJ (NIH), and the results are presented as bar graphs. Error bars indicate the standard error of the mean (SEM). Statistical significance was assessed using Student’s *t*-test: *p* < 0.05 (*), *p* < 0.01 (**). Non-significant differences are indicated as “ns”.

In regard to Figure 5B, we conducted control experiments to rule out RNA contamination in our genomic DNA (gDNA) samples (Figure S4A and B). As shown in Figure S4A, a small amount of RNA contamination was detected in gDNA extracted from BME26 cells or eggs (arrows). This contamination was effectively eliminated by RNase A treatment (Figure S4A, lanes 3 and 6). The purified gDNA samples exhibited high levels of ^6^mA methylation (Figure S4B).

Similarly, in reference to Figure S5A-D, control experiments were performed to confirm that the detected m^5^C methylation (Figure 5C) was specific to gDNA. In this case, no m^5^C methylation was observed in RNA (RNA is present, as shown in Panels A and C) from either BME26 cells or eggs (Figure S5A-D).

### Biological effects of epigenetic inhibitors on BME cells

BME26 cells were tested with three epigenetic inhibitors (Figure 6A, B, C, E,H) that have been extensively validated in mammalian systems: STM2457, a synthetic inhibitor of METTL3^66^; 5’-azacytidine (5’-AZA), an inhibitor of DNA methyltransferases (DNMTs)^67^ ; and Trichostatin A (TSA), an inhibitor of histone deacetylases (HDACs)^68^.

**Figure 6.**
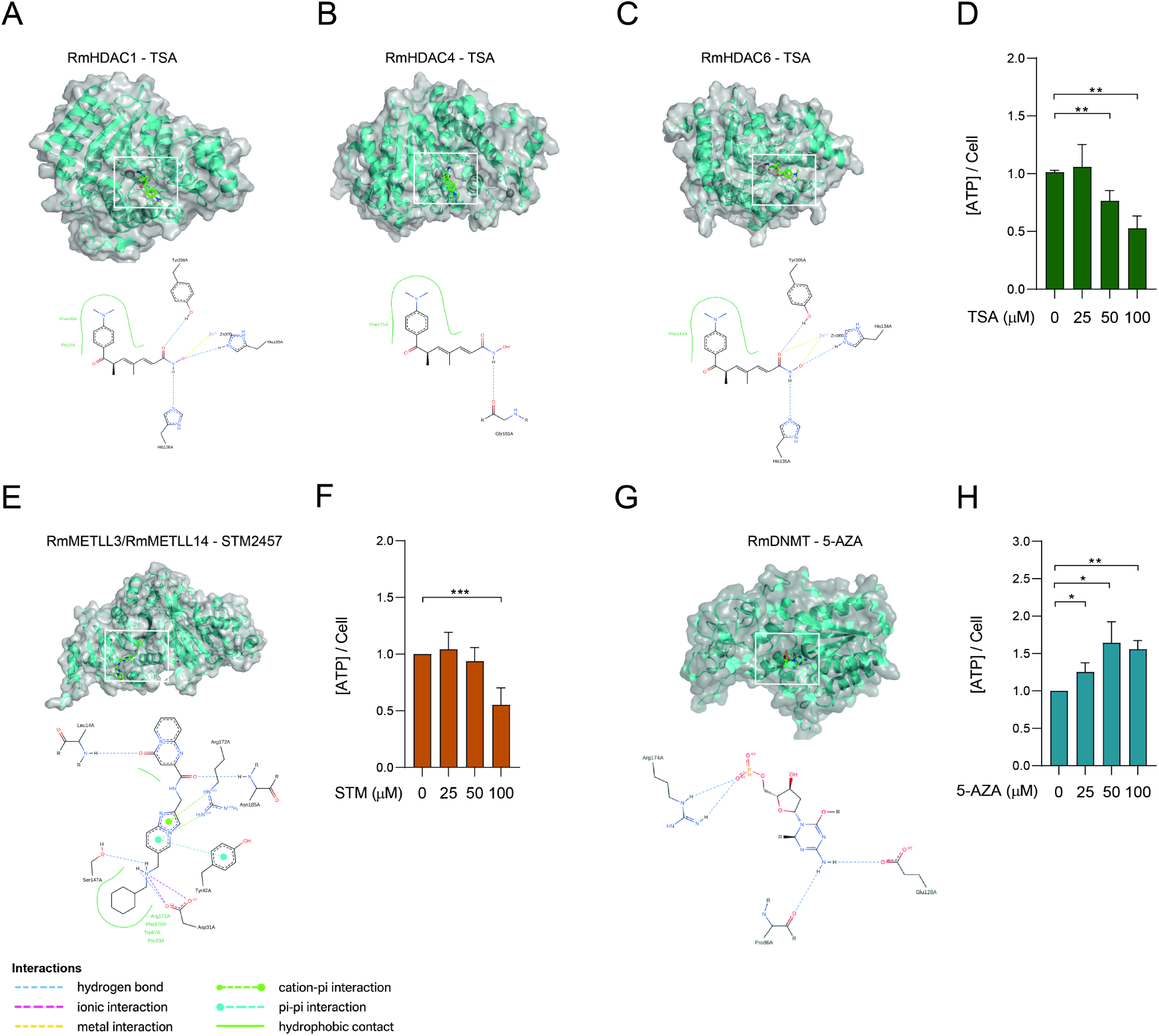
Molecular docking of epigenetic inhibitors to their respective target enzymes. (A-C) Trichostatin A (TSA; green sticks) docked into the substrate tunnel of the histone deacetylases RmHDAC1, RmHDAC4, RmHDAC6. The hydroxamate zinc-chelating moiety coordinates the catalytic Zn²⁺ ion, while the aromatic “cap” group occupies the enzyme surface groove. Hydrogen bonds to the conserved active site residues are shown as blue dashed lines. (E) Ribbon diagram of the METLL3/METLL14-like heterodimer (α-subunit and β-subunit), in complex with STM2457 (green sticks), docked into the conserved SAM-binding pocket. Key hydrogen bonds (dashed lines) and hydrophobic contacts are highlighted. (G) Docking of 5-azacytidine (5’-AZA) into the active site of DNA methyltransferase RmDNMT residues (dashed lines), and π–π stacking with a neighboring aromatic nucleotides. All panels are rendered in PyMOL; water molecules and non-essential ions were omitted for clarity. (D,F,H) BME26 were incubated with increasing concentrations of TSA, STM2457 or 5’-AZA, respectively, for 48 hours cell viability was evaluated by measuring ATP rates. Error bars indicate the standard error of the mean (SEM). Statistical significance was assessed using Student’s *t*-test: *p* < 0.05 (*), *p* < 0.01 (**).

Prior to treatment, we performed molecular docking simulations to evaluate the potential binding of these compounds to their respective tick homologs (Figure 6A-H). Specifically, TSA was docked with HDAC1, HDAC4, and HDAC6, representing Class I, Class IIa, and Class IIb HDACs, respectively (Figure 6A-C); STM2457 was docked with the METTL3/METLL14 heterodimer (Figure 6E) and 5’-AZA was docked with DNMT1 (Figure 6G). The docking simulations revealed that all compounds engage their tick enzyme targets with very favorable binding energies and in virtually identical poses to those observed in the corresponding human complexes (Figure 6, panels A–C, E and G). TSA docks into the catalytic tunnel of tick HDAC1, HDAC4, and HDAC6 with calculated binding free energies of –10.1 kcal/mol, –9.1 kcal/mol, and –10.5 kcal/mol, respectively. In each case, the hydroxamate zinc**-**chelating group of TSA coordinates the active**-**site Zn²⁺ ion in a bidentate fashion, while the cap group forms a network of hydrogen bonds to conserved residues (e.g. Y306 and H141 in HDAC1). A superposition of the tick and human HDAC6•TSA complex (PDB: 5EDU) shows an RMSD of 0.6 Å over all heavy atoms and an identical arrangement of the aromatic “cap” ring against the enzyme surface. Similarly, 5′`-`AZA occupies the DNMT1 catalytic pocket with a binding energy of –8.7 kcal/mol. The nucleoside analogue establishes two key hydrogen bonds between its ring nitrogen and Glu128, and a covalent bond with the side chain of Cys88. The tick**-**DNMT1•5′`-`AZA and human**-**DNMT1•5′`-`AZA structures overlay with an RMSD of under 1 Å, and retain precisely the same hydrogen**-**bonding pattern. Finally, STM2457 binds the methyltransferase heterodimer METTL3–METTL14 with highly favorable energies (–9.3 kcal/mol). Its adenosine**-**mimetic scaffold slots into the SAM**-**binding cleft, forming hydrogen bonds to Asp13 and Asn185 of METTL3, while the imidazo[1,2-a]pyridin-2-yl]methyl] ring tail engages in a conserved stacking interaction with Arg172. Finally, Asp31 and Ser147 interact with the N6-of the [(cyclohexylmethylamino)methyl] moiety. Overlaying the tick and human METTL3 structures confirms an identical binding mode, with both complexes displaying superimposable ligand orientations and contacting residues (RMSD ≈ 0.8 Å). Together, these results demonstrate not only that TSA, 5′`-`AZA, and STM2457 bind strongly to their tick enzyme homologs, but also that their molecular recognition profiles are essentially indistinguishable from those in the human targets.

To assess whether the inhibitors exert cytotoxic effects on BME26 cells, we conducted a cell viability assay based on ATP quantification (Figure 6D, F, and H).

Treatment with either TSA or STM2457 resulted in a significant reduction in ATP levels at concentrations of 50 and 100 μM (Figure 6D and F), indicating decreased cell viability. In contrast, cells treated with 5’-AZA exhibited increased ATP production across the 25 to 100 μM concentration range (Figure 6H), suggesting a potential enhancement of cellular metabolic activity, such as increased mitochondrial function or proliferation. However, the possibility of glycolytic pathway overactivation cannot be ruled out.

To gain more detailed and precise insights into the cytotoxic effects of the inhibitors on cell morphology, we performed confocal microscopy analyses. Looking at BME26 cells treated with TSA for 48h, a clear and significant increase in histone acetylation (H3K27ac) was observed (Figure S6A). We further confirmed histone hyperacetylation in TSA-treated cells by Western blot analysis (Figure S6B). Notably, the Western blot also revealed insights into the crosstalk between H3K27ac and H3K27me3 in BME26 cells, with the acetylation mark (H3K27ac) being overridden by the repressive methylation mark (H3K27me3) (Figure S5B).

In addition to TSA, BME26 cells were also treated with STM2457 and their actin cytoskeleton was analyzed using phalloidin staining (Figure 7). Importantly, the inhibition of RNA-m^6^A methylation in BME26 cells revealed significant alterations in actin filaments (Figure 7A and B, arrows). STM2457-treated cells showed an increase in actin filaments (F-actin) suggesting that RNA-m^6^A methylation is involved in the regulation of actin polymerization in BME26 tick cells. In DMSO-treated cells most of the phalloidin staining was concentrated in small aggregates of F-actin, possibly containing short actin filaments.

**Figure 7.**
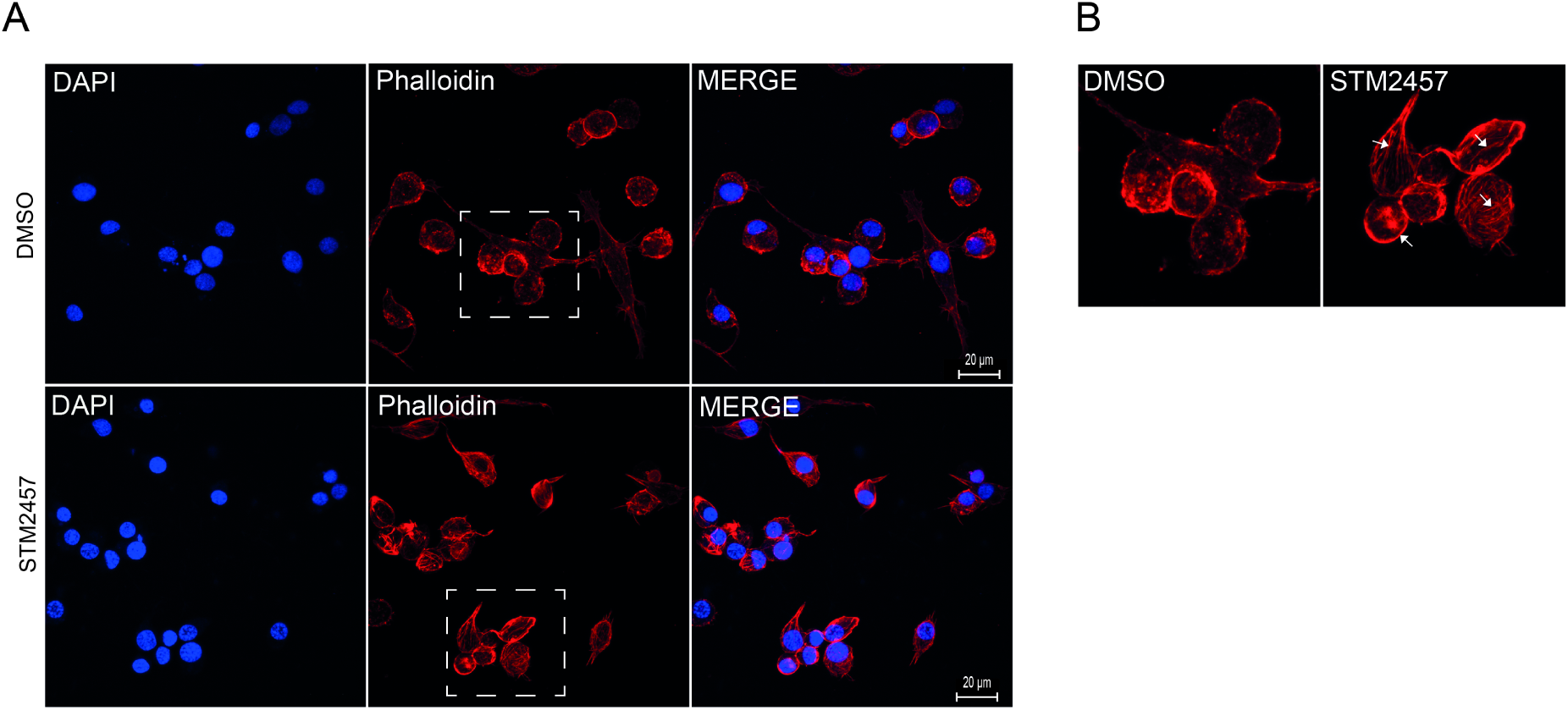
Effect of m^6^A-RNA methylation inhibition on *Rhipicephalus microplus* cells. (A) BME26 cells were treated with either 100 μM STM2457 or DMSO (control) for 48 hours. Nuclei were stained with DAPI, and actin filaments were visualized using Phalloidin. (B) The region highlighted by the square in panel A is enlarged to facilitate visualization of actin fiber organization. Arrows indicate elongated actin filaments observed in cells treated with STM2457.

### m^5^C methylation of the mitochondrial genome and electron transport chain function in BME cells

Following the unexpected rise in intracellular ATP levels upon inhibition of DNA methylation (Figure 6H), we hypothesized that this metabolic shift might be linked to changes in mitochondrial epigenetic regulation—specifically, methylation of the mitochondrial genome. To explore this possibility, we conducted confocal microscopy analyses on BME26 tick embryonic cells treated with 5-AZA. Our observations revealed a marked reduction in m^5^C signals outside the nuclear compartment (Figure 8A), suggesting that mitochondrial DNA may indeed be a target of demethylation under these conditions.

**Figure 8.**
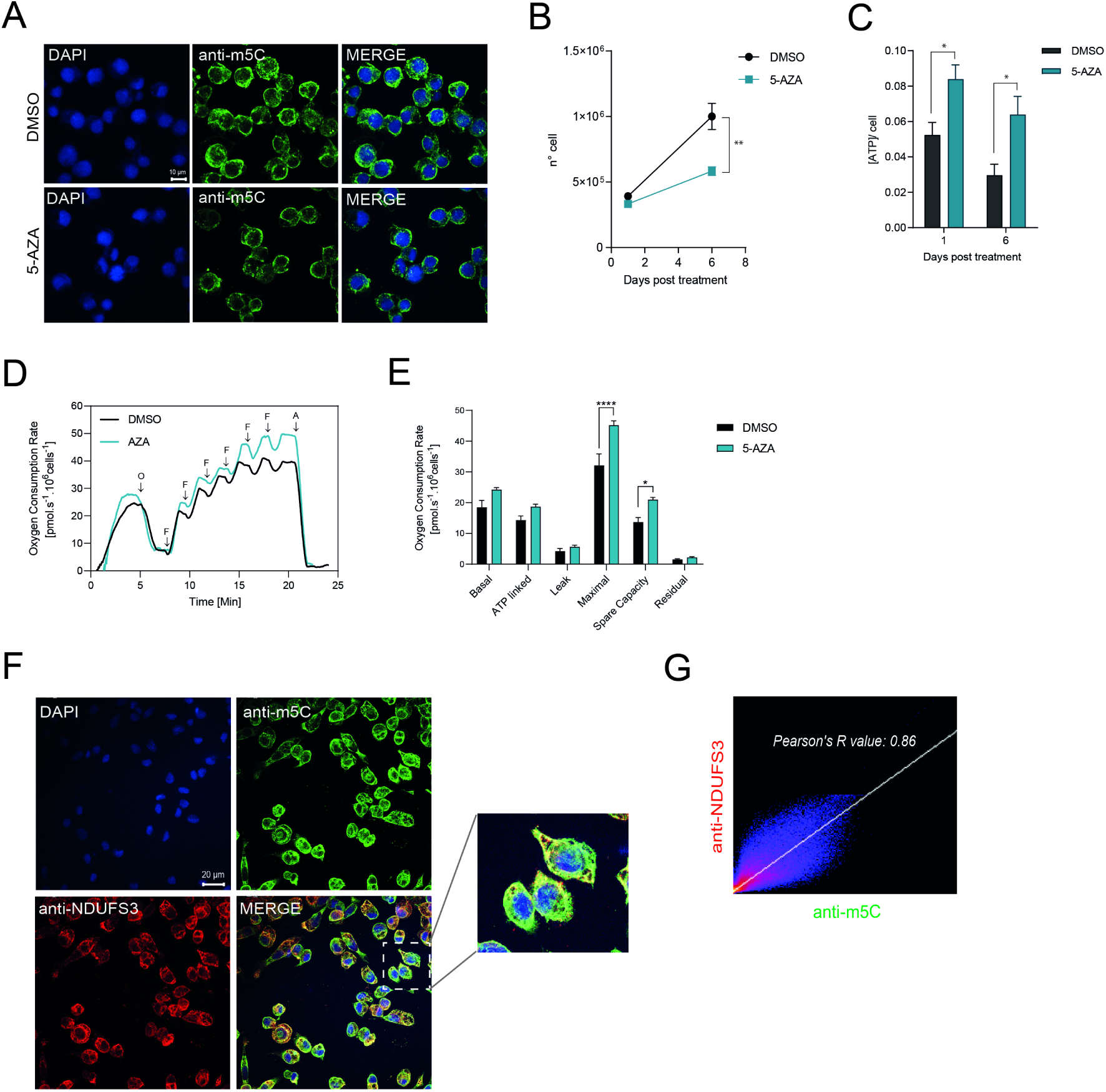
m^5^C-DNA methylation in the mitochondrial genome of *Rhipicephalus microplus*. (A) BME26 cells were treated with 100 μM 5-azacytidine (5’-AZA) for 48 hours to suppress DNA methylation, followed by immunostaining with an anti-m⁵C monoclonal antibody (green) and nuclear counterstaining with DAPI (blue). Confocal microscopy reveals reduced m⁵C signal in 5’-AZA-treated cells compared to control confirming effective inhibition of methylation. Scale bar: 10 μm. (B) The number of cells was counted 6 days post 5’-AZA treatment using a Neubauer chamber. (C) BME26 was incubated with 50 μM of 5’-AZA for six days and cell viability was evaluated by measuring ATP rates. (D) High-resolution respirometry of BME26 cells treated or not with AZA. Representative oxygen consumption trace. O: oligomycin; F: FCCP; A: antimycin A. (E) Mitochondrial respiratory parameters derived from the respirometry data. ****p < 0.0001; *p = 0.0156. *N* = 3, independent cell cultures. Bar graph showing the different components of mitochondrial respiration under various conditions. The parameters include: Basal respiration: The oxygen consumption rate (OCR) under normal, unstimulated conditions, reflecting the energetic demand of the cell at rest. ATP-linked respiration: The portion of OCR directly coupled to ATP synthesis, indicating mitochondrial activity dedicated to energy production. Proton leak (Leak respiration): The residual OCR not coupled to ATP synthesis, representing protons that re-enter the mitochondrial matrix without contributing to ATP production. Maximal respiration: The maximum OCR achieved after uncoupling mitochondrial oxidative phosphorylation, reflecting the total respiratory capacity of the cell. Spare respiratory capacity: The difference between maximal and basal respiration, indicating the cell’s ability to respond to increased energy demand or stress. Non-mitochondrial respiration (Residual): The OCR remaining after inhibition of mitochondrial respiration, representing oxygen consumption by other cellular processes. (F) BME26 cells were incubated with a monoclonal antibody against m^5^C (green) and a monoclonal antibody targeting NDUFS3, a subunit of Complex I in the mitochondrial respiratory chain (red). Nuclei were stained with DAPI (blue). The square region highlights the merged images showing the overlap between NDUFS3 and m^5^C DNA methylation signals. (G) Pearson’s correlation coefficient (R-value) was calculated, demonstrating a positive correlation between the m^5^C signal and mitochondrial localization.

Furthermore, over the course of six days post-treatment with 5’-AZA, we noted a progressive and statistically significant decrease in the total number of viable cells (Figure 8B). Intriguingly, this reduction in cell number was consistently accompanied by a sustained and significant elevation in ATP levels (Figure 8C). These findings point toward a potential link between mitochondrial DNA methylation status and bioenergetic homeostasis in BME26 cells. To confirm this, we performed respirometry experiments and clearly demonstrated that cellular respiration rates were significantly elevated in cells treated with 5’-AZA, as shown in Figure 8D and E. This increase in oxygen consumption strongly suggests an overactivation of the mitochondrial electron transport chain, implying that DNA demethylation induced by 5’-AZA may enhance mitochondrial metabolic activity.

Lastly, to confirm that m^5^C methylation was present in mitochondrial DNA, we conducted confocal microscopy using a monoclonal antibody against NADH dehydrogenase [ubiquinone] iron-sulfur protein 3 (NDUFS3)—a Complex I component of the electron transport chain—alongside an anti-m^5^C monoclonal antibody (Figure 8F). Merged images revealed elevated levels of m^5^C methylation in mitochondrial DNA compared to nuclear DNA (Figure 8F and inset). A strong positive correlation between m^5^C signal and mitochondrial localization was further confirmed (Figure 8G).

## DISCUSSION

This study provides a detailed characterization of the epigenetic machinery involved in *R. microplus* embryogenesis and offers new insights into how epigenetic regulation may influence tick development and physiology. Our findings significantly expand the current understanding of tick epigenetics^49–51^, a relatively underexplored field in vector biology, and suggest new avenues for potential control strategies through epigenetic modulation.

We identified a full repertoire of conserved epigenetic enzymes in the *R. microplus* genome, including key histone acetyltransferases (HATs), histone methyltransferases (HMTs), histone deacetylases (HDACs), and the enzymatic machinery required for DNA (^5^mC and ^6^mA) and RNA (m^6^A) methylation. High sequence similarities within catalytic domains between tick and mammalian orthologs implies functional conservation and provides a strong rationale for investigating small-molecule inhibitors validated in other systems.

By mapping transcriptional profiles and enzymatic activities during embryonic development, we demonstrate that the expression and function of epigenetic regulators in *R. microplus* are temporally dynamic and stage-specific. For example, the acetyltransferase CBP/p300 shows peak activity during mid-embryogenesis (days 9–12), corresponding to periods of rapid tissue differentiation, while Gcn5 activity increases during late embryogenesis (days 15–21), potentially supporting terminal maturation processes. In contrast, several repressive HMTs (e.g., EZH2 and EHMT) displayed increased activity only during the later stages, suggesting roles in developmental gene silencing and structural condensation.

EZH2 (Enhancer of Zeste Homolog 2), a core component of the Polycomb Repressive Complex 2 (PRC2), is a histone methyltransferase enzyme that plays a crucial role in gene silencing by adding methyl groups to histone H3 at lysine 27^70,71^. EzH2 is essential for early embryonic development, particularly for cell lineage determination, embryonic stem cell (ESC) maintenance, and proper differentiation of trophoblast cells^70–73^. In *Drosophila*, Polycomb group (PcG) proteins are crucial for maintaining the silenced state of genes, particularly homeotic genes, which determine body segment identity^71^.

Histone H3 lysine 27 (H3K27) is an antagonistic switch between PcG-mediated repression and activation by Trithorax group (TrxG) proteins^74–76^. TrxGs counteract PcGs through the H3K27 demethylase UTX and by recruiting the histone acetyltransferase (HAT) CREB-binding protein (CBP) and its paralog p300 to acetylate H3K27 (H3K27ac) and mutually exclude H3K27me^374–76^. Importantly, previous studies have shown that a global loss of H3K27 methylation leads to aberrant accumulation of H3K27 acetylation, driven by the histone acetyltransferases CBP and p300^74^. This hyperacetylation occurs at genomic regions where H3K27 methylation is normally present, including Polycomb group (PcG)-bound promoters and non-lineage-specific enhancer elements^76^.

In this regard, our findings in *R. microplus* are particularly significant. We confirmed that the dynamic interplay between H3K27 acetylation (H3K27ac) and trimethylation (H3K27me^3^) also occurs in this species (Compare Figure 4A and D), suggesting that a similar epigenetic regulatory mechanism is conserved. Given the importance of these modifications in controlling gene expression during development, this interplay could serve as a novel target for strategies aimed at disrupting tick development and reproduction, providing a potential avenue for controlling this economically important ectoparasite. 5-methylcytosine (m^5^C) is the most abundant type of DNA modification in the genomes of diverse species^77^. Previous studies have suggested that the dynamic regulation of m^5^C plays important roles in regulating chromatin architecture and gene expression during mammalian development^78^. Interestingly, m^5^C-DNA methylation is absent in *Aedes aegypti*^23^ and *Drosophila*^79^, suggesting that this epigenetic modification is not universally essential for embryonic development in eukaryotes.

In *R. microplus*, we observed distinct expression patterns among the DNA methyltransferases. DNMT1 was primarily expressed in early embryogenesis (days 6-12), consistent with their roles in *de novo* DNA methylation^80^. Interestingly, DNMT3a and DNMT3b remained highly expressed and active in essentially all stages, suggesting a sustained role in embryonic DNA methylation dynamics in ticks, potentially also including the regulation of repetitive elements or non-coding regions^81,82^.

Recent studies suggest that N^6^-methyladenine (^6^mA) DNA methylation is associated with gene expression across a range of eukaryotic species, including algae, worms, flies, and mammals, indicating a potential epigenetic role for ^6^mA in eukaryotic development^83–85^. However, the mechanisms by which ^6^mA-related epigenetic marks are established and interpreted to regulate gene expression programs remain largely unknown.

Previous research has shown that ^6^mA is highly dynamic during early embryogenesis in *Drosophila*. Interestingly, the temporal pattern of ^6^mA dynamics closely coincides with the maternal-to-zygotic transition (MZT)^86,87^. The Forkhead box (Fox) family protein Jumu acts as a maternal transcription factor by preferentially binding to ^6^mA-marked DNA, thereby regulating embryonic gene expression and contributing to MZT, at least in part through the regulation of Zelda^88^.

In contrast, our results demonstrate that ^6^mA DNA methylation levels remain constant throughout *R. microplus* embryogenesis, suggesting that this epigenetic modification may serve a different role during tick embryonic development.

In *R. microplus*, the m^6^A RNA methylation machinery, specifically the methyltransferase complex components METTL3 and METTL14, was found to be expressed across all developmental stages. However, their enzymatic activity peaked during early embryogenesis, indicating a dynamic shift in post-transcriptional gene regulation during this critical period. Notably, the highest levels of METTL3/14 activity were observed as early as six days post-oviposition, a timeframe that coincides with major developmental milestones. This temporal pattern suggests that m^6^A RNA methylation may play a pivotal role in orchestrating the maternal-to-zygotic transition (MZT), a tightly regulated process in which control of gene expression shifts from maternally deposited transcripts to zygotic genome activation. The enrichment of METTL3/14 activity at this stage implies that m^6^A modifications may be essential for regulating transcript stability, splicing, translation efficiency, or degradation during MZT, thereby contributing to proper embryonic development in this tick species. In fact, deletion of the m^6^A-binding protein YTHDF2 in zebrafish embryos leads to reduction of the decay of m^6^A-modified maternal mRNAs and suppression of the zygotic genome activation. Upon deletion of YTHDF2, the embryos fail to initiate timely MZT, undergo cell-cycle pause, and remain developmentally delayed throughout larval life^89^. In addition, studies have shown that m^6^A methylation regulates the embryonic development of insects through affecting maternal mRNA decay, allowing normal embryogenesis in *Drosophila*^90^.

Pharmacological inhibition of epigenetic regulators in BME26 cells corroborated their functional importance. Histone deacetylase inhibition by TSA led to histone hyperacetylation and a reduction in ATP levels, consistent with disrupted chromatin organization, which decreased cell viability. These effects were mirrored by inhibition of m^6^A-RNA methylation, which also reduced viability and caused visible cytoskeletal disruptions (Figure 7).

Although not yet directly shown in ticks, m^6^A RNA methylation could likely modulate actin filament dynamics indirectly by controlling the expression of mRNAs encoding actin regulators^91,92^. This would be especially relevant during early development, feeding, molting, and immune responses. In this context, key indirect m^6^A-regulated pathways are emerging based on what is known in arthropods and other invertebrates^93,94^.

A striking finding of this study is the elevated level of m^5^C-DNA methylation in mitochondrial DNA (mtDNA) compared to nuclear DNA, visualized via immunofluorescence and supported by confocal colocalization with mitochondrial Complex I marker NDUFS3 (Figure 8). Moreover, inhibition of DNA methylation with 5’-AZA, surprisingly elicited pronounced effects on mitochondrial function, increasing both electron transport chain activity and ATP production. This observation is particularly intriguing given the significantly higher levels of DNA methylation detected in the mitochondrial genome. Our data suggests a functional link between methylation status and mitochondrial bioenergetics in tick embryonic cells.

Mitochondrial remodeling during the mammalian peri-implantation stage represents a critical hallmark event required for successful embryogenesis. During this period, mitochondria undergo functional and structural adaptations to meet the escalating energy demands of the developing embryo. A key component of this remodeling is the upregulation of oxidative phosphorylation (OXPHOS), which plays a pivotal role in sustaining the metabolic needs of post-implantation development. However, this metabolic shift also results in increased production of mitochondrial reactive oxygen species (ROS), elevating oxidative stress and placing mtDNA integrity at risk^95^. Such oxidative damage, if left unchecked, can compromise mtDNA stability and ultimately jeopardize embryonic viability^96,97^.

In line with this concept, our data shows that BME26 tick cells treated with 5-5’-AZA, a DNA methylation inhibitor, exhibited elevated respiratory activity, indicative of enhanced mitochondrial function. However, these same cells also demonstrated a marked reduction in proliferation rates, suggesting a potential trade-off between increased mitochondrial respiration and cellular growth. This observation may reflect a stress response to heightened oxidative metabolism, potentially impairing cell cycle progression or activating damage-response pathways.

These findings open new perspectives on the role of organellar epigenetics in arthropods. Given that mitochondrial function is central to embryonic development and stress responses, targeting mitochondrial methylation machinery might represent a novel and selective strategy for disrupting tick viability.

## CONCLUSION

Our integrative approach reveals that epigenetic regulation is not only present but also highly dynamic, and functionally significant during *R. microplus* embryogenesis.

We identified a complete set of conserved epigenetic enzymes, with their expression and enzymatic activities varying during development, suggesting precise control of gene expression and chromatin state throughout embryogenesis.

Importantly, the dynamic interplay between H3K27 acetylation and methylation, suggests a conserved Polycomb/Trithorax regulation in ticks. We also showed that m^6^A-RNA methylation peaks during early embryogenesis, coinciding with the maternal-to-zygotic transition (MZT), supporting a role in post-transcriptional control of gene expression.

Notably, we detected high levels of m^5^C in mitochondrial DNA, and inhibition of DNA methylation leads to a significant overactivation of the mitochondrial electron transport chain. This suggests a previously unrecognized link between mitochondrial DNA methylation and energy metabolism in tick embryonic cells.

Altogether, our results reveal that both nuclear and mitochondrial epigenetic regulation are critical for tick embryonic development. These mechanisms may represent promising targets for the development of novel, epigenetic-based tick control strategies.

## ACKNOWLEDGMENTS

We thank Dr. Marcus Fernandes de Oliveira (UFRJ) for supplying antibody against NDUFS3. This work was supported by the Conselho Nacional de Desenvolvimento Científico e Tecnológico, CNPq (grant number 401737/2023-3 and 443294/2024-0), Fundação de Amparo à Pesquisa do Estado do Rio de Janeiro, Faperj (grant number SEI-260003/001743/2023) and Instituto Nacional de Ciência e Tecnologia em Entomologia Molecular, INCT-EM (grant number 573959/2008-0). LT was supported by the Division of Intramural Research of the National Institute of Allergy and Infectious Diseases (Z01 AI001337-01).

## AUTHOR CONTRIBUTIONS

Conceptualization, MRF, AMA and CL; Methodology, BM, AFA, MFO, SR, SHJ, LFP, MU; Software, JDPLY, LT, ISVJ; Formal analysis, AG, AP, DR, RNF, TMV, CSM; Investigation; AMA, DMO, MPNCS, MFO, SR; Data Curation, AMA, MFO, AG and MRF; Writing - Original draft, AMA and MRF; Writing – Review, all authors; Supervision, MRF; Funding Acquisition, MRF, CL, ISVJ. Project Administration, AMA and MRF.

---

**Supplemental Figure 1.** Protein sequence alignment of full-length epigenetic regulators from *Rhipicephalus microplus* (Rm), human (Hs), and bovine (Bt). Amino acids highlighted in grey represent conserved residues, while those in black indicate identical residues across species.

**Supplemental Figure 2.** (A,B). Phylogenetic trees using Ezh and CBP-p300 orthologs across different species, with domain architecture representation. The tree on the left represents evolutionary relationships based on sequence similarity. Bootstrap support values are indicated at each node. On the right, colored bars represent conserved protein domains, as annotated in the legend. The highlighted sequence from *Rhipicephalus microplus* indicates our species of interest. Domain annotations are based on Pfam.

**Supplemental Figure 3.** SDS-PAGE and Western blot analysis of total protein extracts from *Rhipicephalus microplus* eggs. Eggs from days 1 (1D) to 21 (21D) were probed for the presence of histones H3 and H4 using monoclonal antibodies. Histone levels are notably absent at days 1 and 3, as indicated by the lack of corresponding bands.

**Supplemental Figure 4.** Detection of m^6^A methylation in genomic DNA of *Rhipicephalus microplus*. (A) Agarose gel electrophoresis of genomic DNA isolated from BME26 cells and eggs. The upper band (arrow) corresponds to genomic DNA, while the lower band (arrow) indicates RNA contamination. Treatment with RNase A effectively removed RNA contaminants (lanes 3 and 6). (B) Genomic DNAs from cells or eggs that were treated with RNase A were evaluated by Dot blot analysis showing the presence of m^6^A methylation in RNA-free genomic DNA.

**Supplemental Figure 5.** Lack of 5-methylcytosine (m⁵C) modification in total RNA isolated from *Rhipicephalus microplus*. (A, C) Agarose gel electrophoresis of total RNA isolated from BME26 cells (A) and embryos collected from day 3 to day 21 (C), confirming RNA integrity. (B, D) Dot blot analysis showing no detectable m^5^C methylation in RNA from BME26 cells (B) or embryos (D). Genomic DNA from the same samples was used as a positive control, demonstrating the presence of m^5^C DNA methylation (gDNA).

**Supplemental Figure 6.** Hyperacetylation of histones in BME26 cells. (A) BME26 cells were treated with 100 μM TSA for 48 hours followed by immunostaining with an anti-H3K27ac monoclonal antibody (green) and nuclear counterstaining with DAPI (blue). Confocal microscopy reveals a significantly higher H3K27 acetylation, compared to control, confirming effective inhibition of HDAC activities. Scale bar: 10 μm. (B). Western blot analysis of histone modifications in BME26 cells (as shown in panel A). Protein extracts were analyzed for specific histone modifications to assess epigenetic changes. Increased levels of acetylation at histone H3 lysine 14 (H3K14ac) and lysine 27 (H3K27ac) indicate hyperacetylation in response to the experimental condition. In contrast, a reduction in trimethylation at H3K27 (H3K27me3) was detected, supporting the antagonistic relationship between acetylation and methylation at this residue.

