## Supplementary figures and images for "Unravelling the role of epigenetic regulators during embryonic development of *Rhipicephalus microplus*"

### Supplemental Figure 1

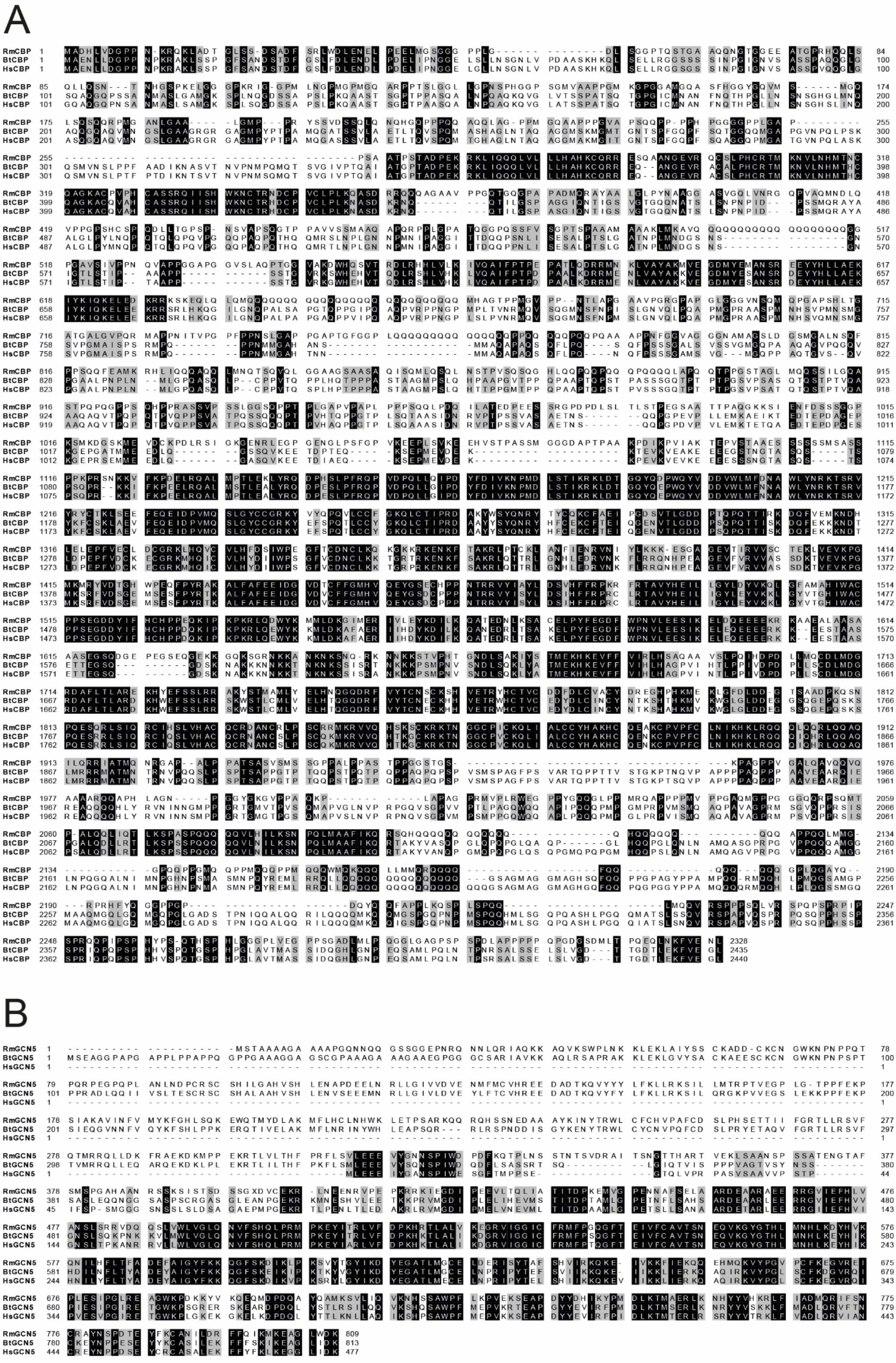

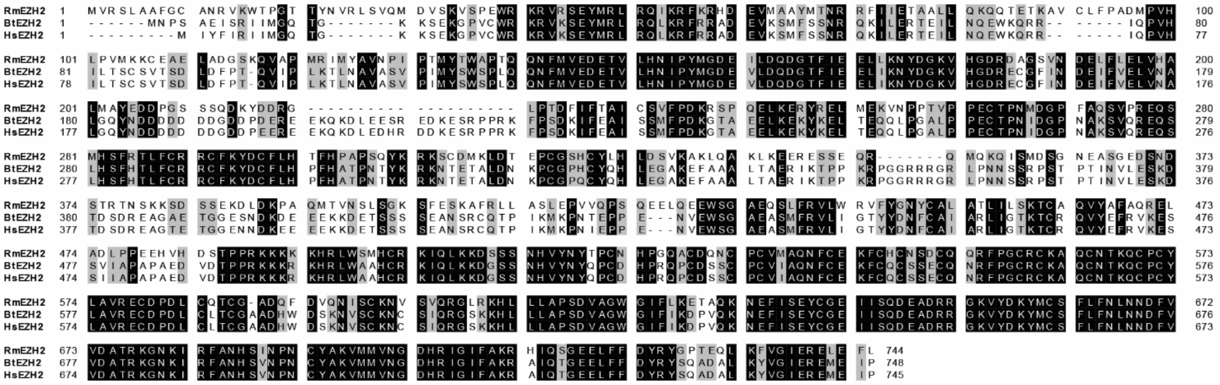

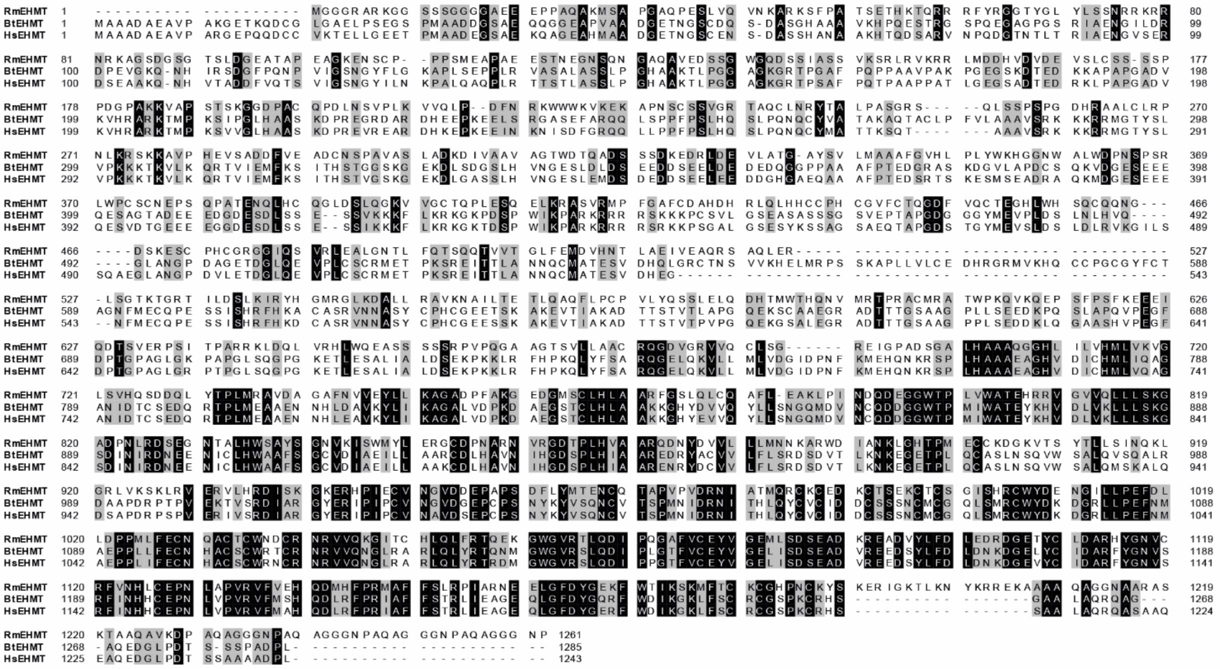

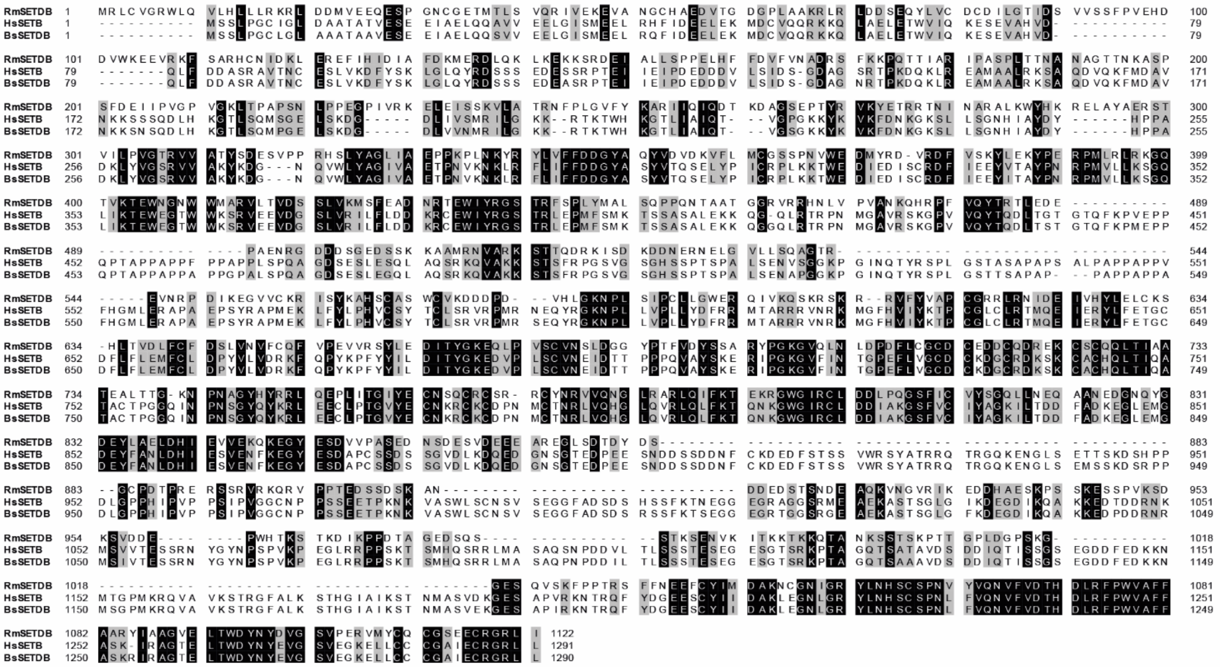

F

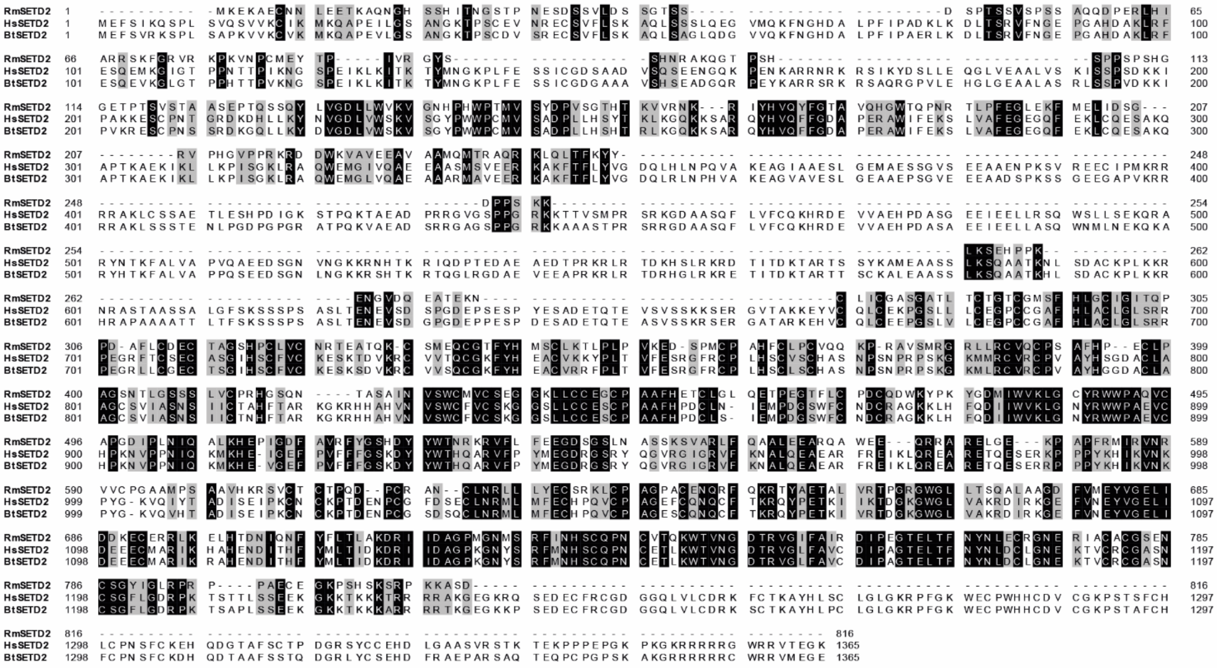

G

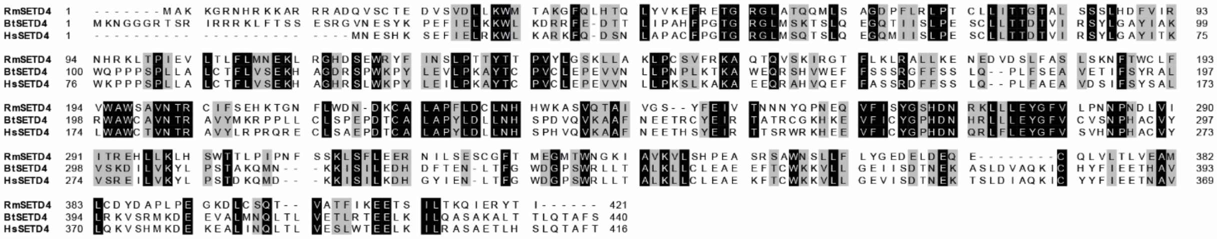

H

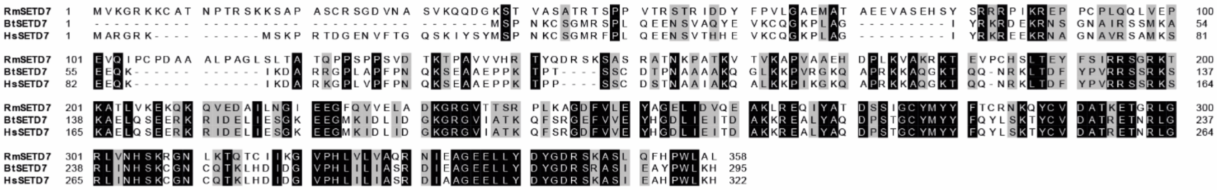

I

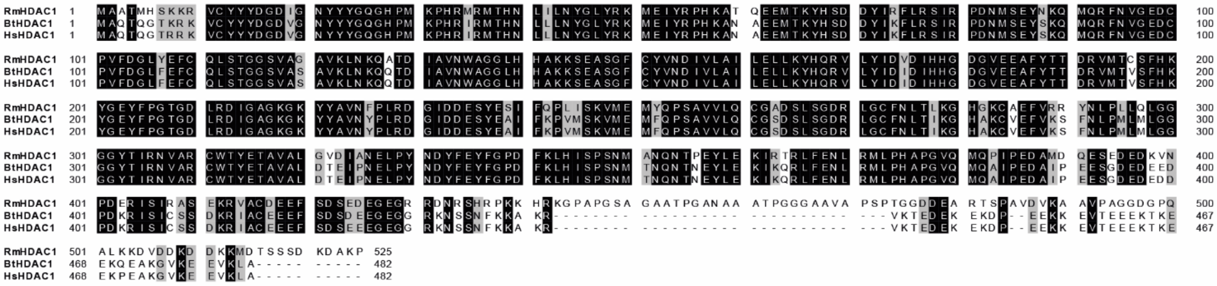

J

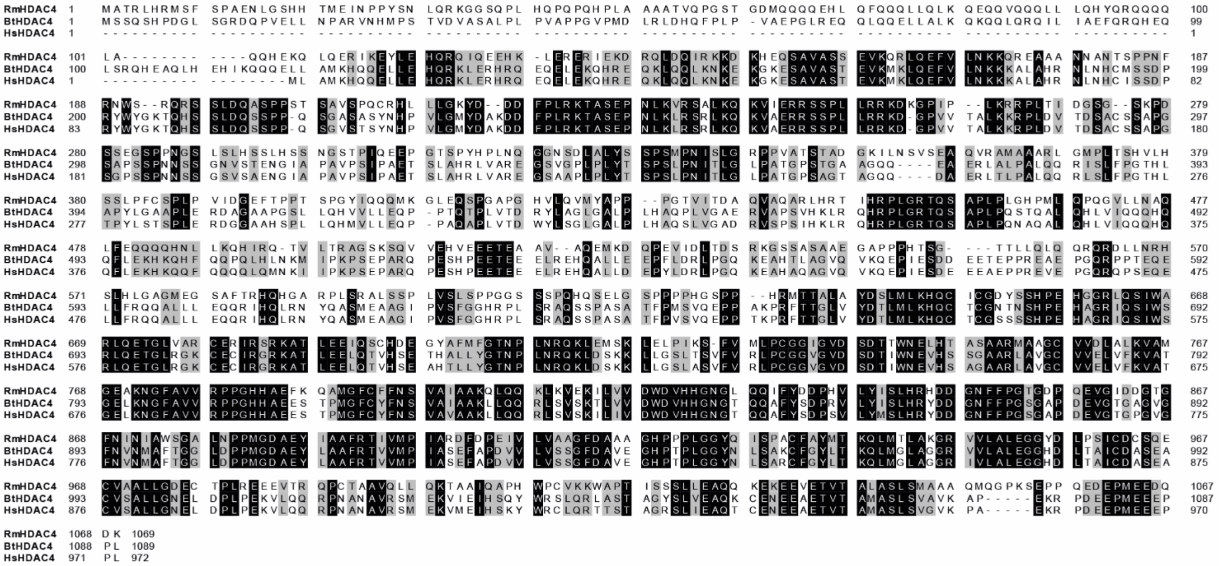

K

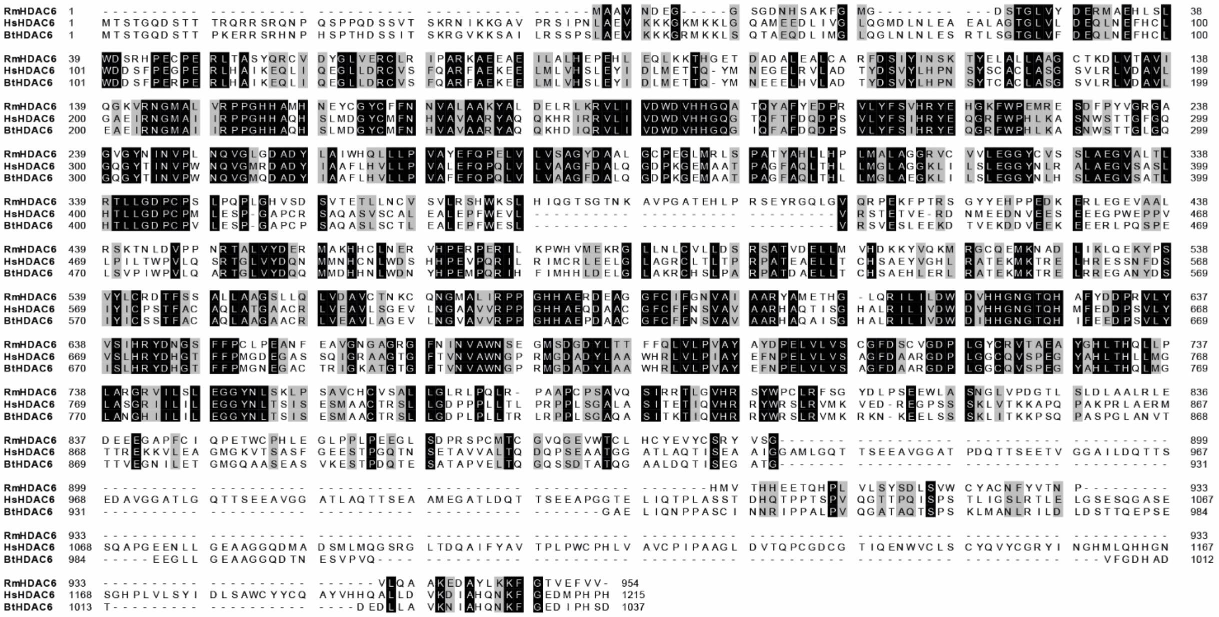

L

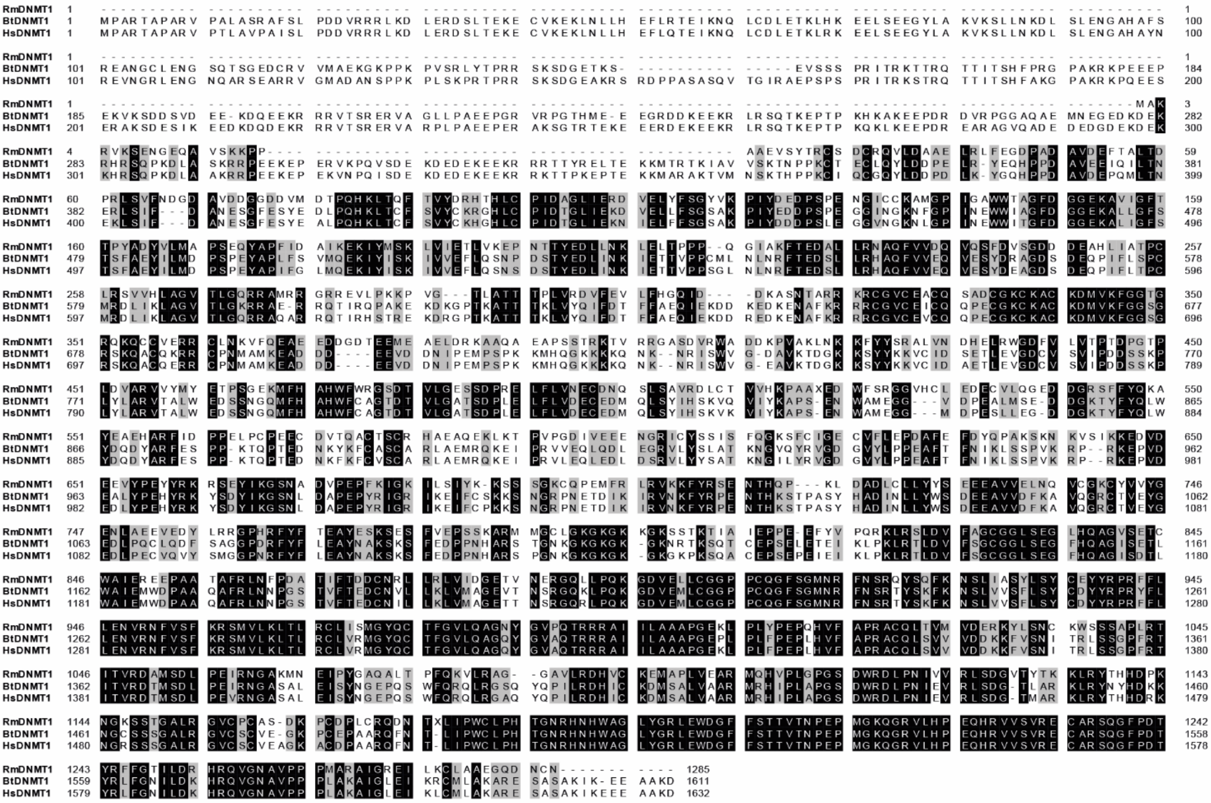

M

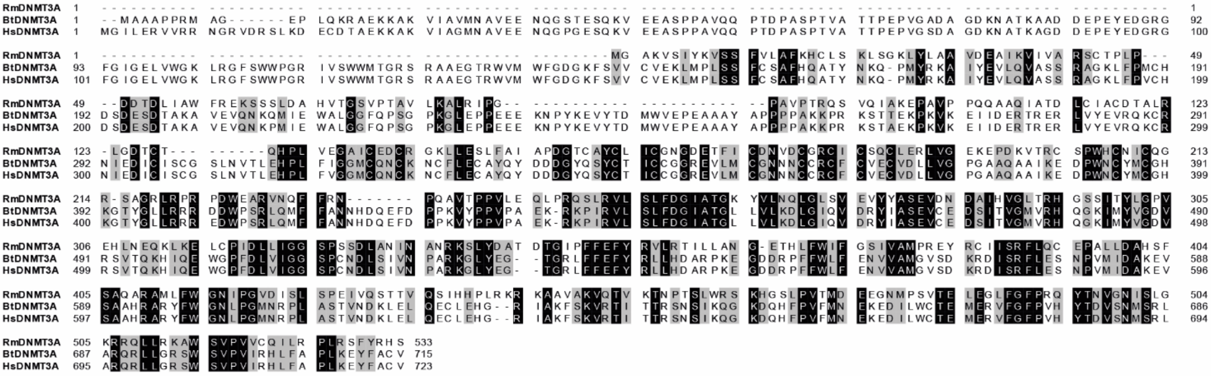

N

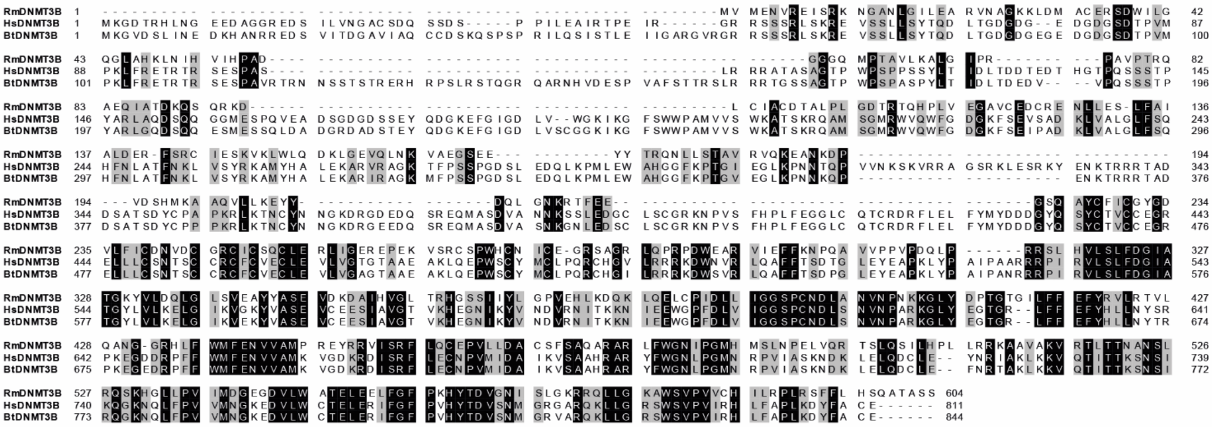

O

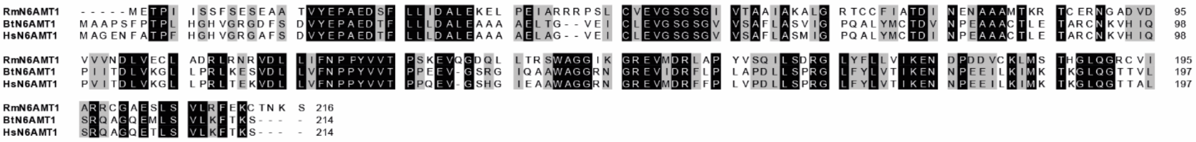

P

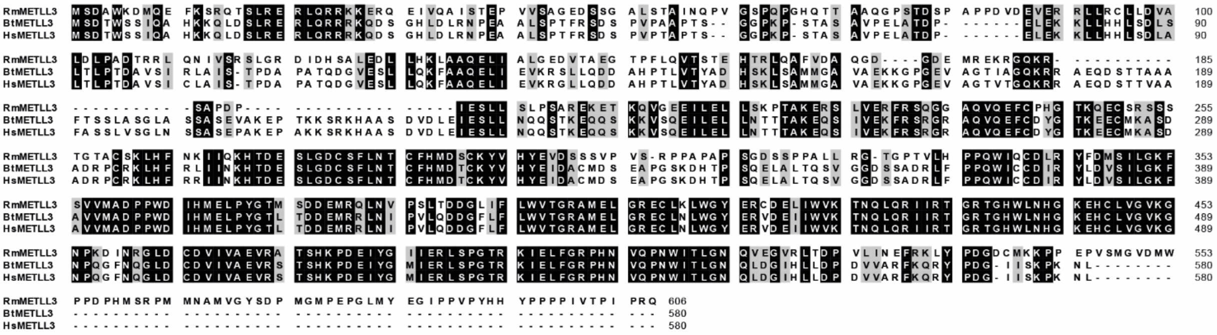

Q

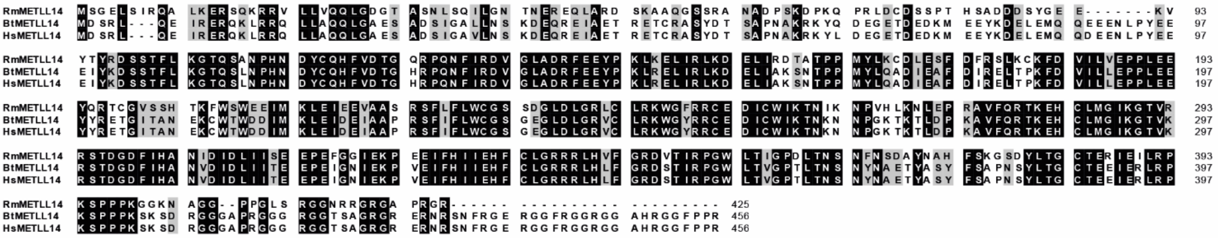
