## Supplemental Figures for "Unravelling the role of epigenetic regulators during embryonic development of *Rhipicephalus microplus*"

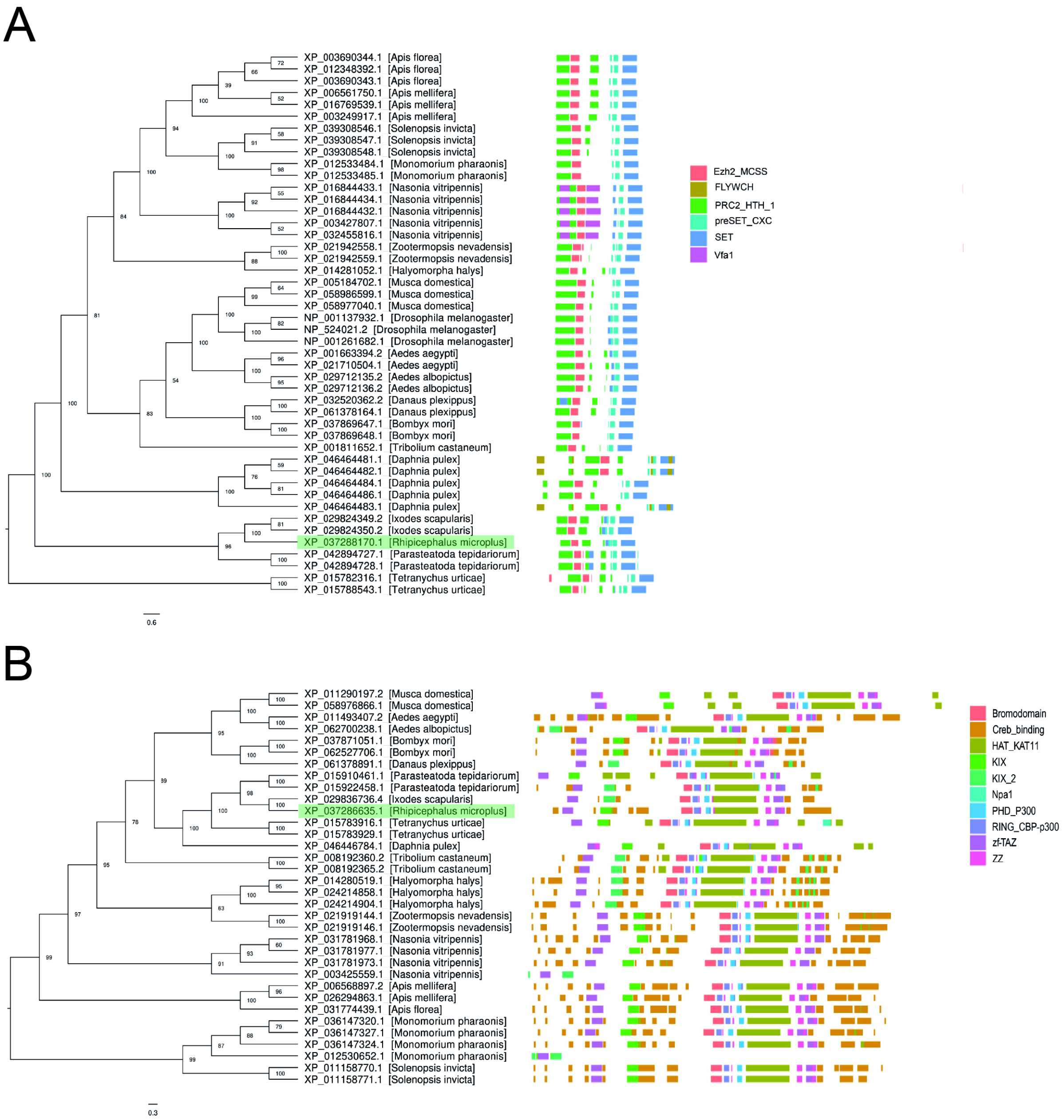


Supplemental Figure 2.

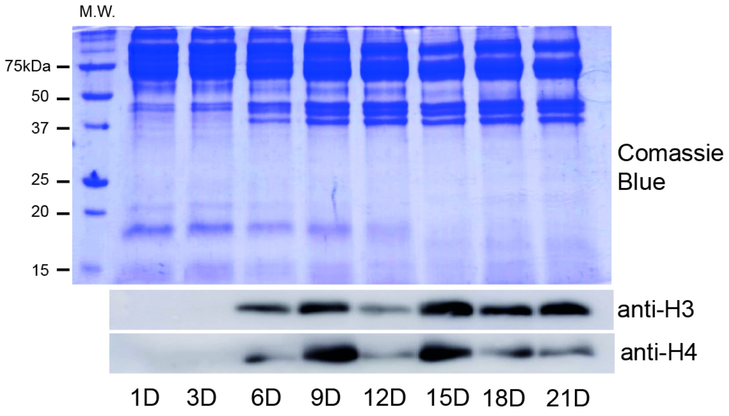


Supplemental Figure 3.

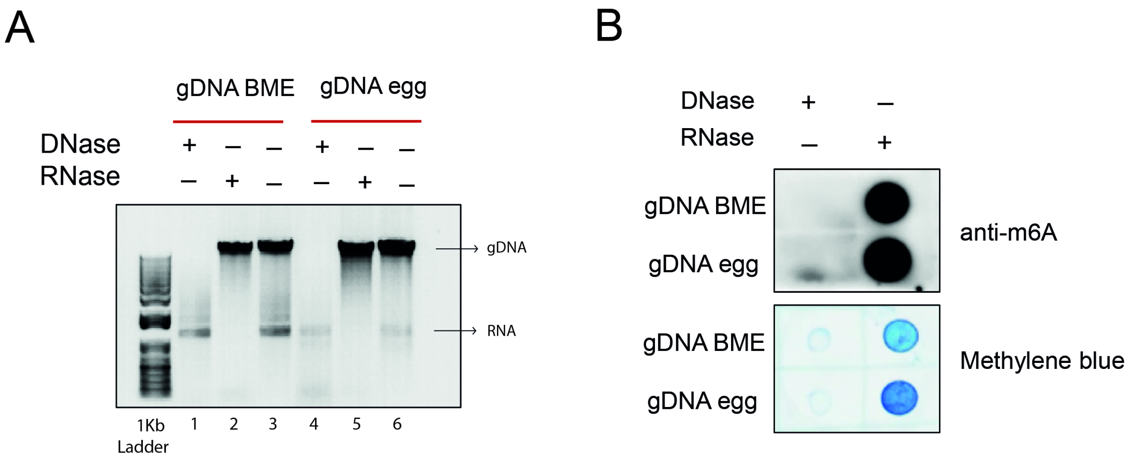


Supplemental Figure 4.

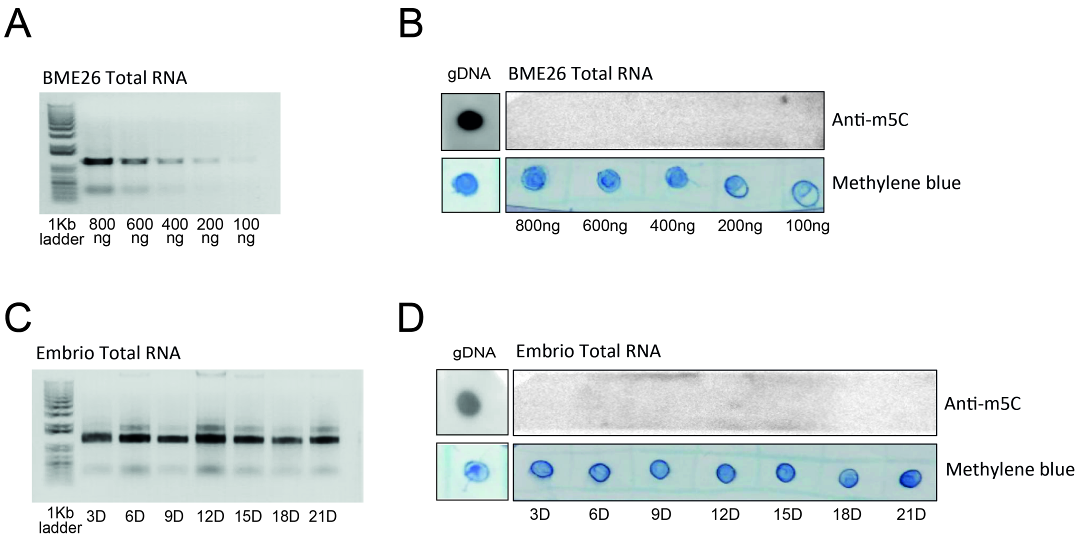


Supplemental Figure 5.

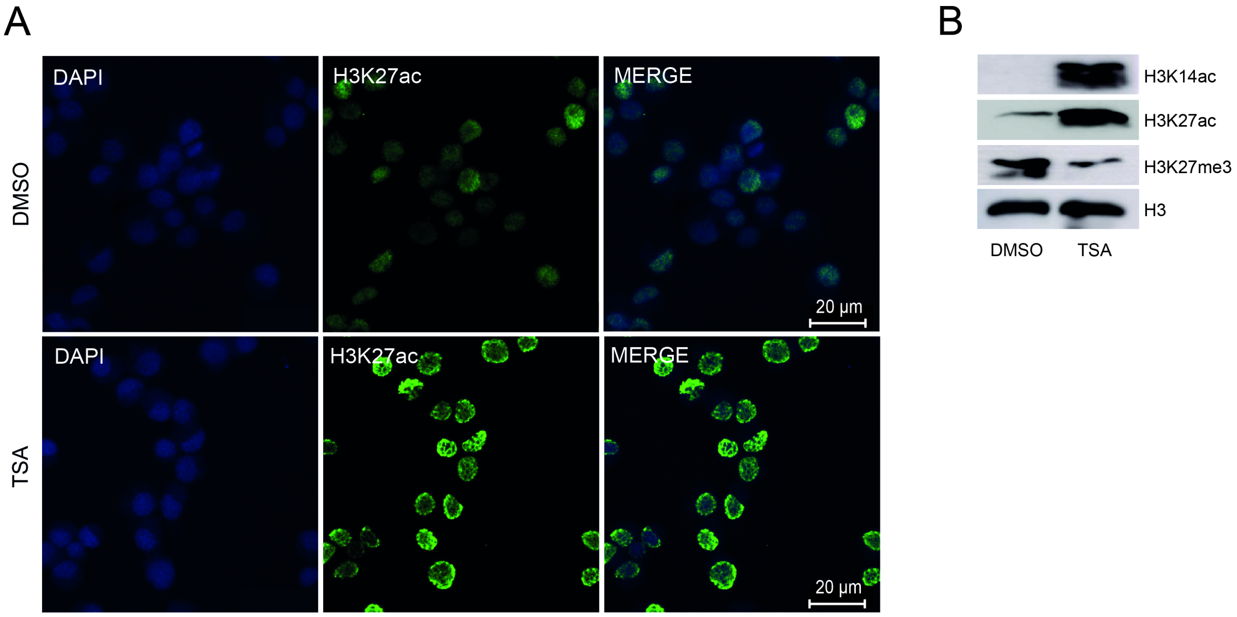


Supplemental Figure 6.

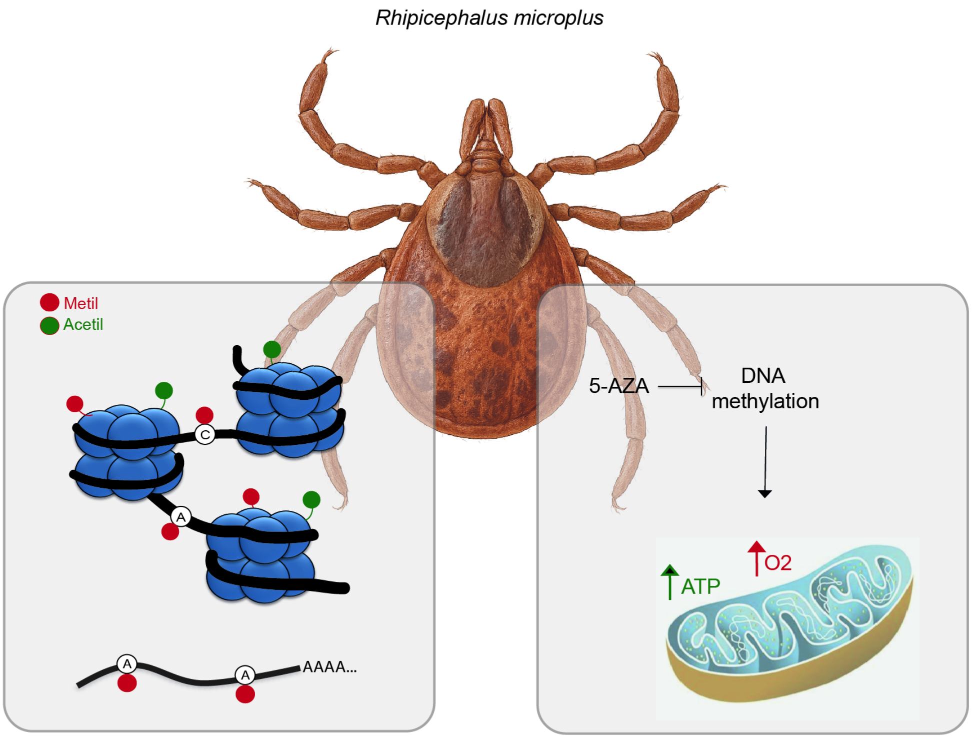


Graphical Abstract
